# SMartini: automated small molecule parametrization for Martini 3 force field

**DOI:** 10.64898/2026.09.29.755481

**Authors:** Danis Yangaliev, S. Banu Ozkan

**Author notes:** Send correspondence to: Danis Yangaliev,.

## Abstract

We present SMartini, an automated pipeline for deriving Martini 3 coarse-grained force-field parameters for arbitrary small molecules. Starting from a molecular structure or a SMILES string, the pipeline maps the all-atom molecule onto a coarse-grained bead representation and automatically fits both bonded and non-bonded parameters. Bonded parameters are obtained via Boltzmann inversion of atomistic molecular dynamics trajectories; successive rounds of coarse-grained simulation and distribution-matching updates then refine these estimates until the coarse-grained conformational ensemble reproduces the all-atom reference within tolerance. We validate the pipeline on a diverse set of small molecules including drug-like compounds, metabolites, and cofactors. Implemented as a modular sequence of scripts that can be orchestrated on high-performance computing clusters, SMartini makes automated parametrization of large ligand libraries practical and reduces the manual effort required for coarse-grained model development, enabling high-throughput coarse-grained simulations of protein–ligand systems within the Martini 3 ecosystem.

**Significance:** The accurate simulation of protein–ligand interactions at physiologically relevant timescales remains a central challenge in computational biophysics and drug discovery. Coarse-grained models offer the necessary computational efficiency, but deriving reliable parameters for small molecules has traditionally been a labor-intensive, expert-driven process that limits throughput. We address this bottleneck by combining a partitioning algorithm with iterative all-atom-to-coarse-grained distribution matching, yielding a fully automated, reproducible pipeline that generates coarse-grained parameters in a fraction of the time required for manual parametrization.

## INTRODUCTION

Protein–ligand interactions underpin virtually all biological processes, from enzymatic catalysis and signal transduction to gene regulation and metabolism. Understanding the thermodynamics and kinetics of ligand binding is therefore central to both fundamental biochemistry and rational drug design. Molecular dynamics (MD) simulations have become an indispensable tool for probing these interactions at atomic resolution. Modern all-atom (AA) force fields—including the CHARMM^1,2^, AMBER^3,4^, and Open Force Field (OpenFF) families^5–7^ — now predict ligand geometries, solvation free energies, and protein–ligand binding affinities with steadily improving accuracy^8^. However, AA simulations remain computationally expensive. Sampling the full binding/ unbinding pathway of a ligand within its protein receptor or larger macromolecular complexes often requires microsecond-to millisecond timescales that are prohibitive for large-scale screening campaigns or complex multi-component systems.

To overcome these limitations, coarse-grained (CG) models reduce the number of degrees of freedom by grouping several heavy atoms into a single interaction site, or bead^9^. The resulting speedup—often two to three orders of magnitude—brings physiologically relevant timescales within reach, making CG molecular dynamics (CGMD) an attractive tool for studying protein– ligand recognition^10^ membrane permeation, and self-assembly processes. Among the many CG schemes available, the Martini force field^11,12^ has emerged as one of the most widely adopted, owing to its systematic parameterization philosophy and broad chemical coverage. Originally developed for lipid membranes^11^, Martini has been successively extended to proteins^13, 14^, carbohydrates^15^, protein–DNA complexes^16^, and RNA^17^. These advances have enabled CG simulations of increasingly complex biomolecular systems, including ribosomes^18–20^, microtubules^21^, viruses^22^ and more. Realizing the full potential of these large-scale simulations, however, also requires well-parameterized small molecules—ligands, cofactors, and metabolites—that interact with the macromolecular components^23^. Recent efforts have produced Martini 3 parameters for ATP^24^, photosynthetic chromophores^25^ and a growing library of drug-like compounds^26^.

Nevertheless, parametrization of arbitrary small molecules remains a largely manual, expert-driven task. For the growing chemical libraries used in virtual screening—often containing hundreds to thousands of compounds—manual parametrization is impractical. Several tools have been developed to automate aspects of this workflow^28^. The AutoMartini M3 pipeline^28^ provides an algorithm for mapping small molecules onto CG beads and assigning Martini 3 bead types. Unfortunately, AutoMartini M3 relies on an exhaustive enumeration of all possible bead partitions, an approach that scales poorly: for molecules beyond roughly 30 heavy atoms (e.g., ATP-sized), the mapping step alone can take hours and consume tens of gigabytes of memory, rendering it impractical for many biologically relevant ligands and for larger cofactors such as heme or chlorophyll^25^. Moreover, automated fitting of bonded parameters—an equally critical step—remains, to our knowledge, even less developed.

Here we present SMartini, a pipeline designed to overcome these limitations, that easily scales to ligands the size of chlorophyll (∼64 heavy atoms) and beyond. We validate SMartini on a diverse set of drug-like molecules, metabolites, and cofactors—including adenosine triphosphate (ATP), chlorophyll, and photosynthetic chromophores —and demonstrate that the pipeline produces numerically stable topologies. Crucially, SMartini automates bonded-parameter derivation through iterative distribution matching against AA MD reference data, substantially reducing the manual fitting effort that limits existing tools. The modular architecture of SMartini allows individual steps to be run independently or orchestrated on high-performance computing clusters, making it practical for parametrizing large compound libraries for downstream CG simulations of protein– ligand systems.

## METHODOLOGY

### General Strategy

The workflow begins with a fragment-based partitioning engine that decomposes the molecule into CG beads while respecting chemical connectivity, ring integrity, and bead-size constraints. Crucially, rather than exhaustively enumerating all mappings, our algorithm solves the partitioning problem locally on chemically meaningful subgraphs and stitches the results together, reducing the combinatorial complexity substantially. Bead types are assigned using per-bead SMILES and logP-based classification, following AutoMartiniM3^28^. For the AA reference, SMartini builds a solvated all-atom system using the OpenFF Sage force field^7^ via the SMIRNOFF parameter assignment scheme5 and runs production MD in OpenMM^29^. Initial bonded parameters are obtained by Boltzmann inversion of the AA trajectory^30,31^. The pipeline then enters an iterative refinement cycle in which CG simulations are run with the current topology and bonded parameters are updated by matching the CG conformational distributions to the AA reference until convergence. To ensure numerical stability, we employ refined angle and dihedral potential forms that mitigate instabilities previously observed in Martini CG models^32,33^.

### Mapping and Bead Type Assignment

The initial stage of the Martini 3 parametrization procedure involves mapping the all-atom structure onto a CG representation by grouping two to six heavy atoms into individual beads. Automated mapping can become the most computationally expensive stage of a coarse-graining workflow. Because the search space expands combinatorially, this global strategy suffers from exponential scaling and becomes impractical for molecules requiring more than roughly 11 beads.

SMartini bypasses this bottleneck entirely by using a purely graph-based partitioning engine that operates locally on the molecular topology. Rather than treating the molecule as a single global system, the algorithm identifies structural building blocks directly from the chemical graph. Individual rings and heavy atoms with a high degree of connectivity are selected as initial fragments. These seeds are then expanded to capture surrounding linear bead segments while keeping functional groups intact to preserve their chemical identity. By solving the mapping problem locally within these functional subgraphs and stitching them together, SMartini reduces the algorithmic complexity from exponential to nearly linear with respect to the number of fragments and heavy atoms. Consequently, systems that are well beyond the reach of exhaustive approaches are partitioned in seconds. For instance, chlorophyll a (CLA, 64 heavy atoms, 27 beads) is mapped by SMartini in under two seconds on a single CPU core, rendering the pipeline highly scalable for large macrocycles and complex cofactors.

Following the mapping stage, bead types are assigned by adopting the chemical typing rules and non-bonded parameter assignments established by the AutoMartini framework^28^. Specifically, each bead is classified by extracting the SMILES string of its constituent atoms and evaluating its partition coefficient (logP) from a reference database. The logP value, together with the number of atoms composing the bead, then determines the final Martini particle classification—polar (P), intermediate (N), nonpolar (C), or charged (D, Q)—as well as its size—regular, small, or tiny.

### Bonded Parameters

The complete set of bonds, angles, and dihedrals is inferred directly from the coarse-grained molecular graph based on bead connectivity. Each bonded parameter is then fitted to target distributions extracted from the automated AA reference trajectory. By default, a 20 ns AA trajectory is collected, which provides well-converged conformational distributions across all tested molecules; in practice, even a 2 ns simulation yields distributions of sufficient quality for reliable bonded-parameter fitting.

A well-known caveat is that unbiased AA simulations sample only the thermally accessible regions of conformational space: free-energy wells separated by barriers higher than a few k_B_Tmay remain unvisited on the simulation timescale. The classic example is alanine dipeptide, where even the two major basins of the Ramachandran map may not be fully explored in a short AA trajectory^34^. This limitation is an intrinsic challenge for any coarse-graining scheme that derives effective potentials from equilibrium AA sampling. While enhanced sampling methods such as metadynamics^35^ or replica exchange MD^36^ can be deployed to overcome these free-energy barriers and map the full conformational landscape, standard equilibrium trajectories remain the practical default for automated setups. Initial bonded parameters are obtained by Boltzmann inversion^30^ of these AA distributions.

Starting from the Boltzmann-inverted parameters, the pipeline refines the bonded terms through an iterative AA-to-CG feedback loop (three iterations by default). At each step, a short CG simulation is performed with the current topology, the resulting distributions are compared to the AA reference, and the parameters are updated to minimize the discrepancy between the two ensembles. In most cases, this procedure converges well within the default three iterations, producing a final CG model that accurately reproduces the main conformational features of the atomistic reference. Convergence can be slower, however, for molecules with strongly coupled angle and dihedral degrees of freedom, a limitation discussed below.

### Molecular Dynamics Simulations

#### All-atom reference simulations

All-atom MD simulations of the solvated ligands were performed with OpenMM 8.5^29^. The ligand was parameterized with the OpenFF 2.1.0 (Sage) force field^7^ via the SMIRNOFF assignment scheme^5^; solvent and ions used the Amber19 force field with OPC water. Each ligand was solvated in a dodecahedral box with 1.2 nm padding, and the ionic strength was set to zero. The system was energy-minimized for up to 1 000 steps, heated gradually from 30 K to 300 K over 10 cycles (1 000 steps per cycle at 1 fs), and then equilibrated for 10 ps in the isothermal-isobaric (NPT) ensemble with a 1 fs timestep and a Monte Carlo barostat at 1 bar. Production runs used a 2 fs timestep with the LangevinMiddleIntegrator (friction coefficient γ = 1.0 ps^−1^) at 300 K and 1 bar. Electrostatic interactions were treated with the particle-mesh Ewald (PME) method and a 1.0 nm cutoff; bonds involving hydrogen were constrained, and water molecules were kept rigid. The default production length was 20 ns (10 M steps); in practice, even 2 ns of AA sampling proved sufficient for reliable bonded parameter extraction, though this does not account for a priori undersampling of the full conformational landscape. Trajectory manipulation, feature extraction, and data preprocessing across the pipeline were performed using MDAnalysis^37,38^.

#### Coarse-grained simulations

All CG simulations were performed with GROMACS 2023^39,40^ using the Martini 3 force field^12^ and standard Martini water (four water molecules per bead). The solute was placed in a cubic box with 1.0 nm minimum distance to the box edge and solvated using a bead radius of 0.17 nm; ions were added to achieve charge neutrality where necessary. The system was energy-minimized using the steepest descent algorithm (force tolerance 100 kJ mol^−1^ nm^−1^), equilibrated for 100 ps in the NPT ensemble, and then subjected to production simulations with a 10 fs integration timestep. The temperature was controlled at 300 K with the velocity-rescale thermostat^41^ (coupling constant *τ* = 1.0 ps), and the pressure was maintained at 1 bar with the c-rescale barostat (*τ* = 4.0 ps, compressibility 3.0 × 10^−4^ bar^−1^, isotropic coupling). Electrostatic interactions were cut off at 1.1 nm using the reaction-field method with a relative permittivity of 15^42^; Lennard-Jones interactions were cut off at 1.1 nm with the potential-shift-Verlet modifier. No bond constraints were applied. Each iterative fitting cycle used a 10 ns CG simulation (1 M steps); three iterations were performed by default (see Results).

Unlike proteins, for which the standard Martini 20 fs timestep is well established, small-molecule ligands are typically significantly stiffer owing to the prevalence of aromatic rings, conjugated double bonds, and compact covalent frameworks. The higher-frequency internal degrees of freedom in such molecules necessitate a shorter timestep for stable integration; while 20 fs can be employed for some systems, it does not generalise reliably across the diverse chemistries encountered in small-molecule libraries. All CG production simulations reported here therefore used a timestep of 10 fs.

## RESULTS

### Validated Molecules

To evaluate the SMartini pipeline, we selected a diverse set of seven small molecules spanning a range of sizes and chemical functionalities (Table 1). The test set includes ANP, a nucleotide building block; CLA, a large tetrapyrrole macrocycle essential for photosynthesis; 5,6-dimethyl-benzimidazole (DMBI), a benzimidazole derivative; fumaric acid thioester analog (FTA), a small polar compound; heme (HEM), the iron-coordinating porphyrin cofactor; lidocaine benzyl analog (LIB), a drug-like molecule with multiple rotatable bonds; and Δ^9^-tetrahydrocannabinol (THC), a pharmacologically active cannabinoid. These molecules range from 20 to 64 heavy atoms (9 to 27 CG beads), covering apolar, polar, and charged chemistries, and include both rigid macrocycles and flexible drug-like scaffolds. The AA and CG representations of all seven molecules are shown in Figure 1.

**Table 1.** Validated molecules with their heavy atom count, CG bead count, and chemical class.

| Ligand | # HAs | # Beads | Chemical Class |
| --- | --- | --- | --- |
| ANP | 31 | 12 | Nucleotide |
| CLA | 64 | 27 | Tetrapyrrole macrocycle |
| DMBI | 20 | 9 | Benzimidazole |
| FTA | 23 | 11 | Thioester |
| HEM | 42 | 18 | Porphyrin cofactor |
| LIB | 40 | 20 | Drug-like |
| THC | 23 | 9 | Cannabinoid |

**Figure 1.**
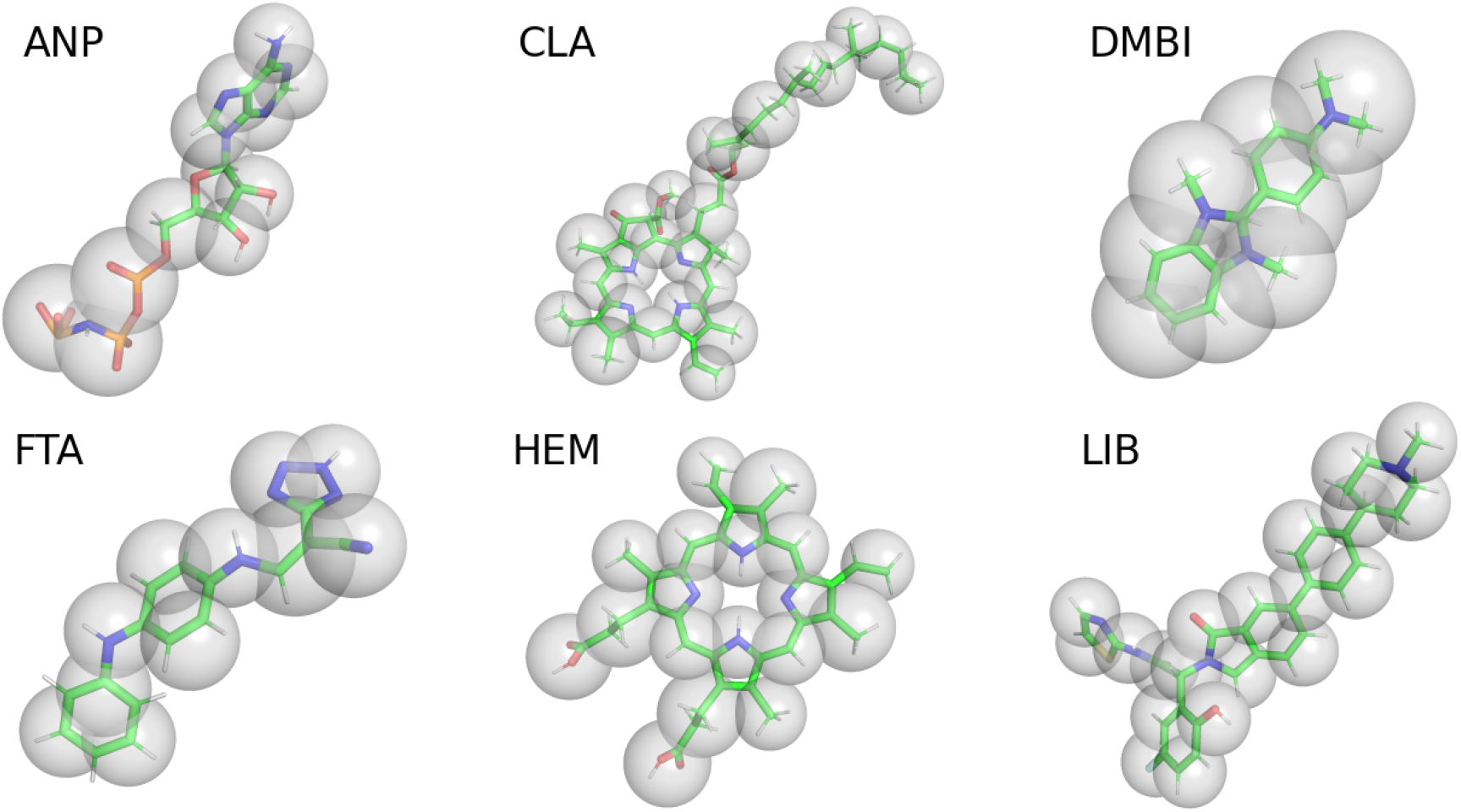
AA (sticks) and CG (transparent spheres) representations of the seven validated molecules. CG beads are shown as semi-transparent spheres overlaid on the all-atom structure; bead sizes correspond to Martini 3 bead-type radii.

For each molecule, the parameterization proceeded without manual intervention: starting from a standard structure data file (SDF), the pipeline automatically partitioned the molecule into CG beads, assigned Martini 3 particle types, generated the all-atom reference trajectory using OpenFF Sage^7^, performed Boltzmann inversion^30^ to obtain initial bonded parameters, and iteratively refined them via AA-to-CG distribution matching.

We note, however, that in contrast to workflows that rely on a global energy function to score and sort all candidate mappings, our approach utilizes targeted heuristic rules. Out of the box, the default pipeline settings comfortably parameterize any arbitrary small molecule. However, because a single global scoring function rarely accounts for every nuance in large or complex structures, localized heuristics give users the option to adjust specific topologies to suit their exact needs. All tunable parameters are exposed in the core configuration file (config.py^43^), with optional flags applied to three distinct cases:

1. LIB: Containing multiple rigid ring assemblies, cross-ring bead fragmentation was explicitly permitted by setting `keep_rings_together=False`.
2. ANP: Virtual sites were enabled (`use_vsites=True`) for the stiff fused rings of adenine base.
3. CLA: Ring symmetrization (`symmetrize_rings=True`) was coupled with explicit user-defined bead specifications (`specify_beads`) to handle the massive, highly symmetrical tetrapyrrole macrocycle properly.

All other molecules in the validation suite were parameterized using the standard default configuration, which serves as the general entry point for any unseen molecule. The complete configuration scripts and output files for all benchmark examples are available in the repository^43^.

### Bonded parameter distributions

To quantitatively assess the agreement between the AA reference and CG conformational ensembles, we computed the normalized 1-Wasserstein distance *W*_1_ = ∫|*F*(*x*) − *G*(*x*)| *dx* for each bonded term category—bonds, angles, and dihedrals—across all validated molecules. This metric measures the similarity of two distributions as the minimal “cost” of transporting the probability mass of one distribution into the shape of the other. Each distance was normalized by the combined data range; values below *∼* 0.05 were deemed satisfactory. Figure 2 presents the per-molecule mean Wasserstein distances, and an example of dihedral distributions for ANP with *W*_1_ = 0.032 to provide visual context for the raw metric values.

**Figure 2.**
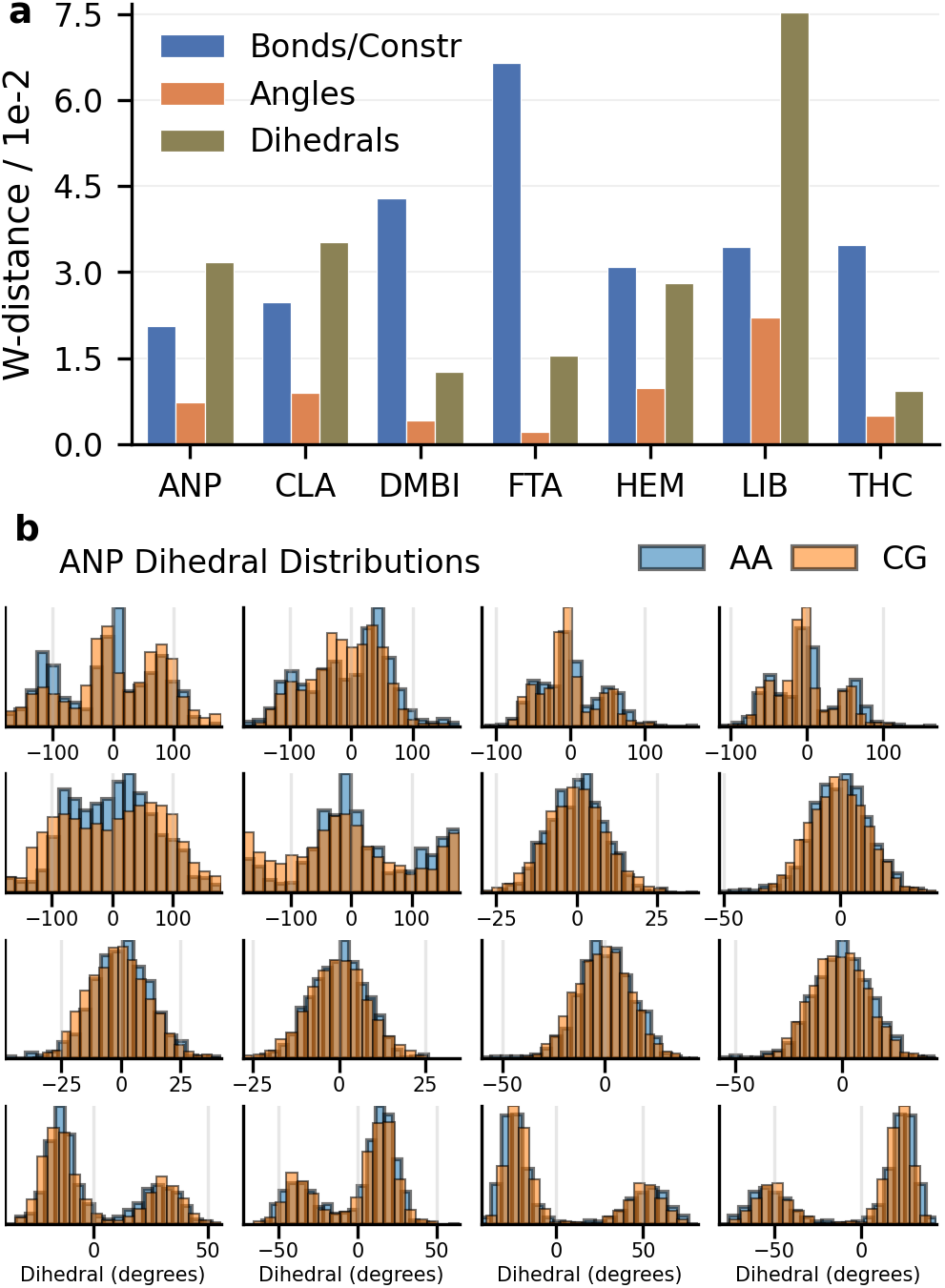
Quality assessment of bonded parameter fitting. (a) Mean normalized Wasserstein distances between AA and CG distributions for bonds/constraints (blue), angles (orange), and dihedrals (green) across all validated molecules. Lower values indicate better agreement. (b) Examples of dihedral distributions for ANP corresponding to a W-distance of 0.032 (see *Supplementary Information* for the full set of distributions at higher resolution).

Across the seven molecules, bond and constraint Wasserstein distances fall predominantly below 0.05, with a mean of 0.036. Angle distances are tightly clustered below 0.01 for six of the seven molecules, with the sole exception of LIB (0.022), which is discussed in detail below. Dihedral distances show somewhat greater variability (0.009–0.075) reflecting the inherent difficulty of fitting periodic torsional potentials with the limited functional forms available in GROMACS. The internal Wasserstein analysis confirms that the iterative distribution-matching procedure employed by SMartini produces CG bonded parameters that are quantitatively suitable for reproducing AA conformational preferences across a chemically diverse ligand set.

Several outlier values merit discussion. The elevated bond Wasserstein distance for FTA (0.067) originates from a mismatch between the stiffness of certain AA bonded degrees of freedom and the corresponding CG constraints. In the all-atom reference, some bonds exhibit very narrow, sharply peaked distributions, whereas the CG topology represents these same degrees of freedom using holonomic constraints that fix the inter-bead distance to a single value. This discrepancy is a necessary trade-off: constraining overly stiff bonds is required for numerical stability under the 10 fs integration step. The observed Wasserstein shift reflects this deliberate replacement of a hard harmonic potential with a rigid constraint, rather than a failure of the fitting procedure.

Small-molecule ligands are typically significantly stiffer than proteins and nucleic acids due to the prevalence of aromatic rings, conjugated bonds, and short-range covalent linkages. While pushing integration increments beyond 10 fs is possible for a subset of rigid molecules, doing so is unreliable across the diverse chemistries encountered in typical ligand libraries. This 10 fs value therefore represents a compromise that ensures numerical stability without sacrificing the computational efficiency gains of coarse-graining. The resulting topologies remain stable in production CG simulations and faithfully reproduce the overall molecular shape and dynamics.

### Solvent-Accessible Surface Area, RMSD, and Radius of Gyration

Beyond internal conformational distributions, we evaluated how well the CG models preserve three global structural descriptors: the solvent-accessible surface area (SASA), the root-mean-square deviation (RMSD) of the CG bead positions relative to the mapped AA reference structure, and the radius of gyration (Rg), which probes the overall compactness of the molecule. SASA probes the exposure of the molecule to the solvent environment and is directly relevant to the quality of protein–ligand binding simulations, while RMSD measures the overall structural drift of the CG ensemble from the AA equilibrium geometry.

Figure 3 presents the mean SASA, RMSD, and Rg values with 95% bootstrap confidence intervals (1 000 replicates, non-parametric; see *Supplementary Information*, Section S2) for both the AA reference trajectories and the CG production runs. SASA was computed using the Shrake– Rupley algorithm^44^ with a probe radius of 0.14 nm as implemented in MDTraj^45^; CG beads were treated as single spheres with their assigned Martini 3 radii (0.17–0.235 nm, Section S3). RMSD was evaluated after optimal rigid-body alignment of each CG frame to the mapped AA reference (Section S4).

**Figure 3.**
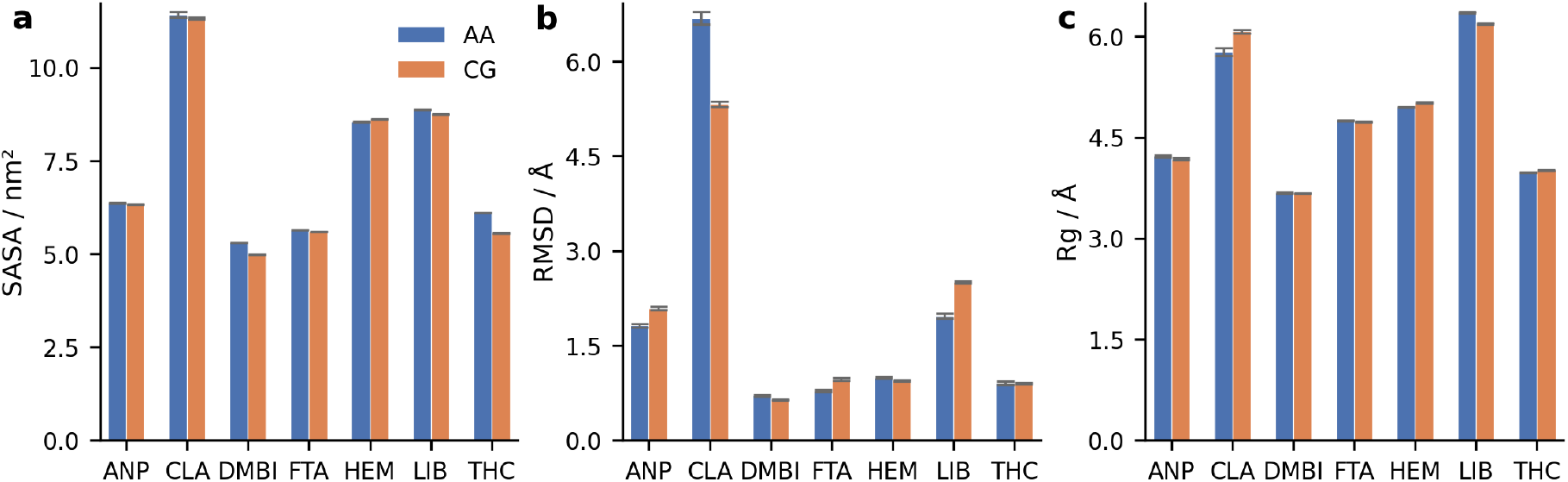
Mean SASA, RMSD, and radius of gyration Rg with 95% bootstrap confidence intervals for AA (blue) and CG (orange) trajectories. (a) CG SASA closely matches AA SASA across all molecules, with only two showing slightly lower CG values, as is typical for Martini 3^25^. (b) RMSD values differ by less than 0.3 Å for five out of seven molecules. (c) Rg values are nearly identical between AA and CG for six molecules.

Across all molecules, the CG SASA closely matches the all-atom, with only two molecules showing slightly lower CG values. Marginally lower SASA values are consistent with previous explicit parametrizations of similar ligands in Martini 3^24,25^. RMSD values are generally comparable between AA and CG; most molecules show differences of less than 0.3 Å, with the exception of two outliers: CLA and LIB. Both compounds proved significantly harder to parameterize using default GROMACS potentials. Specifically, CLA contains a long flexible phytol tail that reduces its numerical stability in the CG representation. Meanwhile, LIB exhibits the second-largest CG RMSD (2.50 Å) and AA–CG RMSD gap (0.5 Å). This deviation is consistent with its elevated angle and dihedral Wasserstein distances. The difficulty of fitting ligands like LIB is discussed in further detail below. The radius of gyration shows near-identical agreement between AA and CG across six of the seven molecules. The sole exception is CLA with a slightly elevated CG Rg relative to the AA reference.

### Protein–Ligand Contact Analysis

To validate SMartini-derived CG topologies in a protein–ligand context, we simulated two protein systems—EGFR kinase domain together with ANP and the HEM-binding protein (PDB ID: 1TQN^46^) —with their respective ligands parametrized by SMartini. For each system, we computed the fraction of native contacts *Q*(*t*)^47^, defined as the fraction of bead–residue pairs within a cutoff *d*_*c*_ = 0.7 nm that remain in contact relative to the reference frame; *Q*(*t*) ranges from 0 (complete loss) to 1 (perfect preservation).

Figure 4 shows the *Q*(*t*) evolution for 1TQN and the kinase domain. In both systems, *Q*(*t*) remains above 0.8 throughout the simulation, and the coarse-grained *Q*(*t*) tracks the all-atom reference reasonably well, indicating that the native binding interface is stably maintained at the CG level. We note, however, that maintaining a ligand stably bound to a protein is not always straightforward in a CG representation, which is a known limitation of CG force fields, for two primary reasons. First, CG models lack explicit hydrogen bonds, which often serve as key anchoring interactions in protein–ligand complexes. Second, the absence of hydrogen atoms can introduce steric mismatches between the AA and CG representations, making it harder to preserve native contacts. In such cases, additional distance restraints that mimic the AA hydrogen-bond network may be required. In the present work, we introduced two supplementary bonds in the kinase domain simulation to maintain the contacts between ANP and its binding pocket.

**Figure 4.**
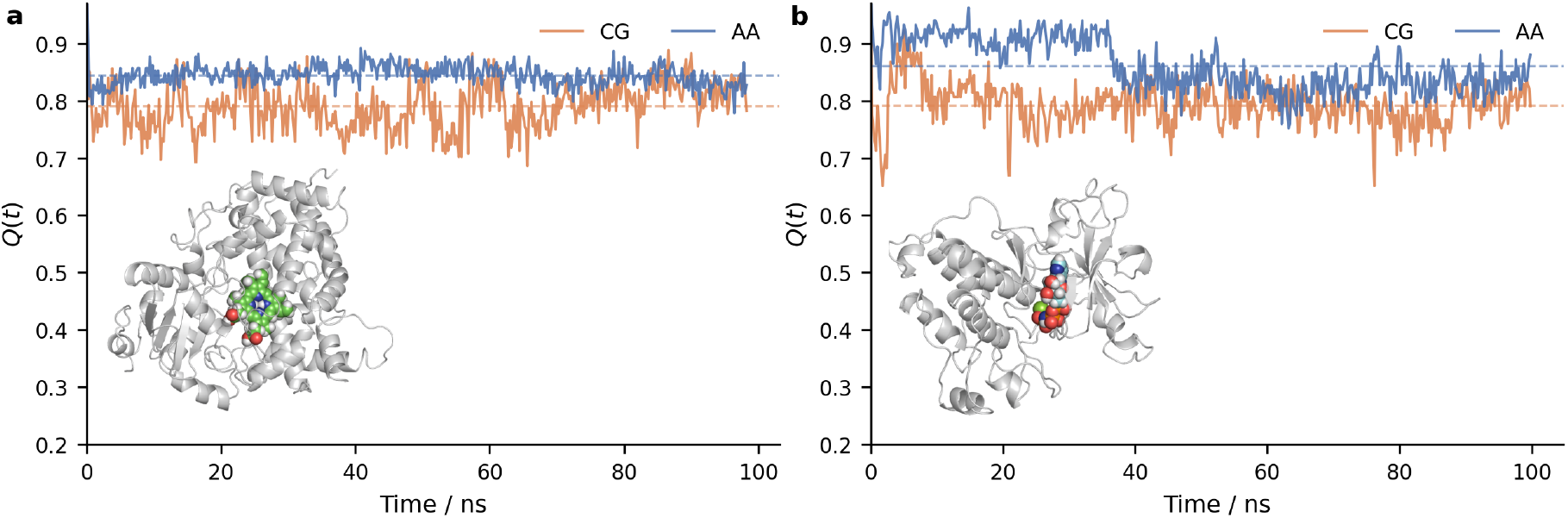
Fraction of native contacts *Q*(*t*) as a function of simulation time for (a) 1TQN and (b) the EGFR kinase domain. In both systems *Q*(*t*) remains above 0.8 throughout the simulation and closely tracks the all-atom reference, indicating that SMartini-generated ligand topologies preserve the native binding interface at the CG level. Two supplementary bonds were introduced in the kinase domain simulation to compensate for the absence of explicit hydrogen bonds in the CG representation.

## DISCUSSION

### Limitations of Bonded-Parameter Fitting

While the overall agreement between AA and CG conformational ensembles is satisfactory, certain bonded terms in flexible, drug-like molecules prove challenging to fit within the constraints of the Martini functional-form library. The most prominent example in our test set is LIB, whose angle W-distance (0.022) and dihedral W-distance (0.075) are markedly higher than those of the other molecules.

The root cause lies in the bonded potentials available in GROMACS: angle interactions are limited to a single-well harmonic form, which cannot represent multimodal distributions. Dihedral angles, by contrast, are fitted with cosine-series potentials that naturally accommodate multiple wells and are generally well behaved in isolation. The difficulty arises when angle and dihedral degrees of freedom are tightly coupled—the harmonic angle potential cannot capture the multimodal character induced by adjacent dihedral substates. In the AA reference, molecules with several rotatable bonds often exhibit intrinsically multimodal dihedral distributions; while the cosine-series fit captures the overall envelope, the corresponding angle distributions become multi-peaked as well, a feature a single-well harmonic cannot reproduce. This effect is particularly pronounced in LIB and CLA, where strong angle–dihedral coupling leads to a complex free-energy surface that cannot be faithfully rendered with the available GROMACS potentials.

These limitations are inherent to the current Martini 3 framework as implemented in GROMACS. A potential route to resolving this problem is to use more flexible simulation engines: the OpenMM implementation of Martini^48^ supports tabulated and custom bonded potentials that could, in principle, capture multi-well angle distributions. We have not yet explored this direction, as the differences in how OpenMM and GROMACS handle constraints would add an additional layer of complexity; nevertheless, it represents a promising avenue for future development. More broadly, introducing tabulated or multi-well bonded potentials into the standard Martini toolchain would directly alleviate these fitting challenges.

## Conclusion

We have presented SMartini, an end-to-end automated pipeline for deriving Martini 3 coarse-grained parameters for arbitrary small molecules. The fragment-based mapping engine handles molecules of 30–60 heavy atoms and beyond, well past the reach of existing tools, and the iterative AA-to-CG distribution matching produces bonded parameters without manual intervention. SMartini reduces expert parametrization from days to minutes, and its modular, HPC-compatible architecture supports deployment in large-scale virtual-screening pipelines.

## ACKNOWLEDGEMENTS

This work was funded by Bill & Melinda Gates Foundation INV-067331 and the National Institutes of Health (grant no. GM147635 to S.B.O.)

## CODE AND DATA AVAILABILITY

The code used in this study is available on GitHub: https://github.com/DanYev/SMartini. Code used in protein-ligand simulations: https://github.com/DanYev/reForge. Molecular Dynamics data are available upon request.

## AUTHOR CONTRIBUTIONS

Conceptualization: D.Y. Formal analysis: D.Y. Funding acquisition: S.B.O. Investigation: D.Y. Methodology: D.Y., S.B.O. Project administration: S.B.O. Writing—original draft: D.Y. Writing— review and editing: D.Y., S.B.O.

