## Supplemental Material for "SMartini: automated small molecule parametrization for Martini 3 force field"

#### S1. Internal Wasserstein Distance

The agreement between all-atom (AA) and coarse-grained (CG) conformational distributions for each bonded term was quantified using the 1-Wasserstein (earth-mover's) distance. For two one-dimensional distributions with cumulative distribution functions $F(x)$ and $G(x)$, the 1-Wasserstein distance is defined as

$$W_1(F, G) = \int_{-\infty}^{\infty} |F(x) - G(x)| \, dx. \tag{1}$$

This metric has a natural geometric interpretation: it measures the minimal "cost" of transporting the probability mass of one distribution into the shape of the other, where the cost is proportional to the distance the mass must be moved. For bonds and constraints, the Wasserstein distance was computed directly on the distance distributions (in nm). For angles, the distributions over $[0, \pi]$ were compared, and for dihedrals, the circular nature of the distributions over $[-\pi, \pi]$ was respected by evaluating the distance on the periodically wrapped cumulative distributions.

To facilitate comparison across terms with different physical units and scales, each Wasserstein distance was normalized by the combined data range of the two distributions:

$$\tilde{W}_1 = \frac{W_1(F, G)}{\max(\text{range}_F, \text{range}_G)}, \tag{2}$$

where $\text{range}_F = \max(F) - \min(F)$. A normalized distance of zero indicates identical distributions; values below approximately 0.05 are considered to reflect satisfactory agreement at CG resolution.

#### S2. Error Estimation (Bootstrap Resampling)

All reported means are accompanied by 95% confidence intervals estimated via bootstrap resampling. For each quantity (SASA, RMSD, Wasserstein distance), the trajectory data were resampled with replacement 1 000 times, and the mean was recomputed for each bootstrap replicate. The lower and upper bounds of the 95% confidence interval were taken as the 2.5th and 97.5th percentiles of the bootstrap distribution, respectively. This non-parametric approach makes no assumptions about the underlying distribution of the data and is robust to the autocorrelation present in MD trajectories because the bootstrap operates at the level of trajectory frames, which are treated as exchangeable observations after adequate equilibration.

#### S3. Solvent-Accessible Surface Area

The solvent-accessible surface area (SASA) was computed using the Shrake–Rupley algorithm (Shrake and Rupley 1973) as implemented in MDTraj (McGibbon et al. 2015). For all-atom trajectories, a probe radius of 0.14 nm was used, corresponding to the approximate radius of a water molecule. For coarse-grained trajectories, each Martini bead was treated as a single sphere with its assigned van der Waals radius (0.17–0.235 nm depending on bead type), and the same 0.14 nm probe radius was employed.

#### S4. Root-Mean-Square Deviation

The root-mean-square deviation (RMSD) of CG bead positions relative to the mapped AA reference was computed as

$$\mathrm{RMSD}(t) = \sqrt{\frac{1}{N}\sum_{i=1}^{N}\left\|\mathbf{r}_i^{\mathrm{CG}}(t) - \mathbf{r}_i^{\mathrm{ref}}\right\|^2}, \tag{3}$$

where $N$ is the number of CG beads, $\mathbf{r}_i^{\mathrm{CG}}(t)$ is the position of bead $i$ at time $t$ in the CG trajectory, and $\mathbf{r}_i^{\mathrm{ref}}$ is the reference position of bead $i$ obtained by mapping the equilibrium AA structure onto the CG representation. The RMSD was computed after optimal rigid-body alignment of each CG frame to the reference structure, using only the CG bead coordinates.

#### S5. Intermolecular Contact Maps and Fraction of Native Contacts

To evaluate how well the CG model reproduces the protein–ligand binding interface observed in atomistic simulations, we compute intermolecular contact maps and the fraction of native contacts $Q(t)$ (Best et al. 2013). A contact is defined between a ligand bead $i$ and a protein residue $j$ when their distance is within a cutoff $d_c = 0.6$ nm, corresponding to the typical Martini nonbonded cutoff range. This bead-to-residue resolution is chosen to make the analysis directly comparable across CG and AA representations: in both cases the protein is reduced to the centre-of-geometry (COG) of each residue, and the ligand coordinates are expressed at the CG-bead level (using the ITP mapping to reduce AA ligand atoms to bead COGs).

For each frame $t$, the binary contact matrix $C_{ij}(t)$ is

$$C_{ij}(t) = \begin{cases} 1, & \text{if } \|\mathbf{r}_i^{\text{lig}}(t) - \mathbf{r}_j^{\text{prot}}(t)\| \leq d_c, \\ 0, & \text{otherwise,} \end{cases} \tag{4}$$

where $\mathbf{r}_i^{\text{lig}}$ is the position of ligand bead $i$ and $\mathbf{r}_j^{\text{prot}}$ is the COG of protein residue $j$. The set of *native contacts* $\mathcal{N}$ is defined as the set of bead–residue pairs in contact in the reference (first) frame of the trajectory:

$$\mathcal{N} = \{(i,j) \mid C_{ij}(0) = 1\}. \tag{5}$$

The fraction of native contacts at time $t$ is then

$$Q(t) = \frac{|\mathcal{N}(t)|}{|\mathcal{N}(0)|}, \quad \text{where } \mathcal{N}(t) = \{(i,j) \in \mathcal{N} \mid C_{ij}(t) = 1\}. \tag{6}$$

$Q(t)$ ranges from 0 (complete loss of the native interface) to 1 (perfect preservation) and provides a simple, interpretable measure of binding-site stability over the course of the simulation.

To focus the analysis on the binding pocket, we apply a proximity filter: only protein residues whose COG lies within 1.0 nm of any ligand bead in the reference frame are retained. This spatial pre-filtering removes residues that are far from the binding site and could otherwise dominate the contact statistics. To ensure that CG and AA contact maps are directly comparable, the residue selection is further unified: only residues that pass the 1.0 nm proximity filter in *both* representations are included in the final contact matrices.

Trajectory I/O and atom selection for CG systems were handled using MDAnalysis (Gowers et al. 2016); all-atom trajectory processing used MDTraj (McGibbon et al. 2015). Per-frame pairwise distance matrices were computed in a vectorised form using NumPy (Harris et al. 2020). Contact frequency maps were visualised as heatmaps with identical axes for CG and AA, enabling a side-by-side comparison of the binding interface sampled by each representation.

### S6. Supporting Figures

#### ANP

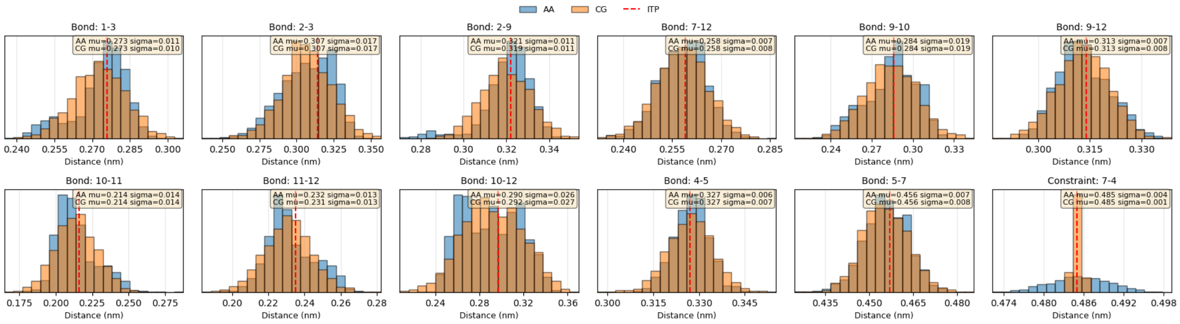


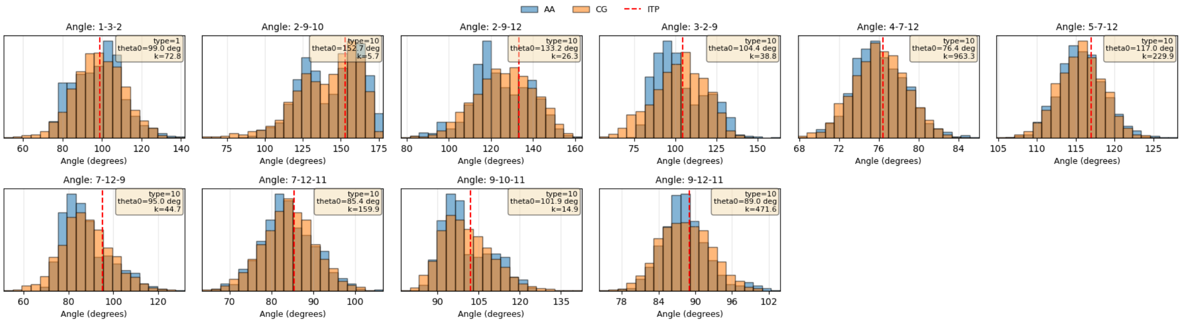


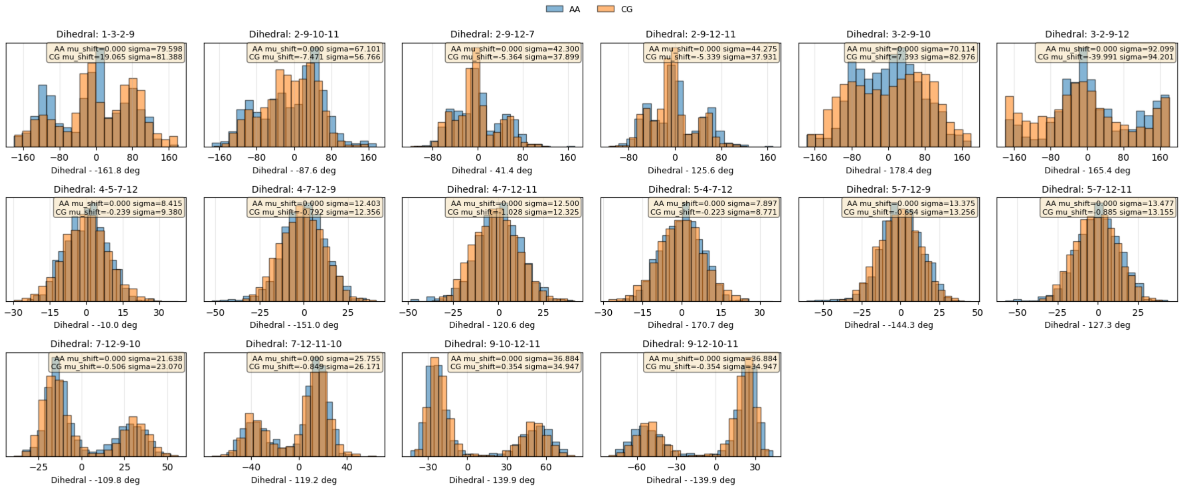

### CLA

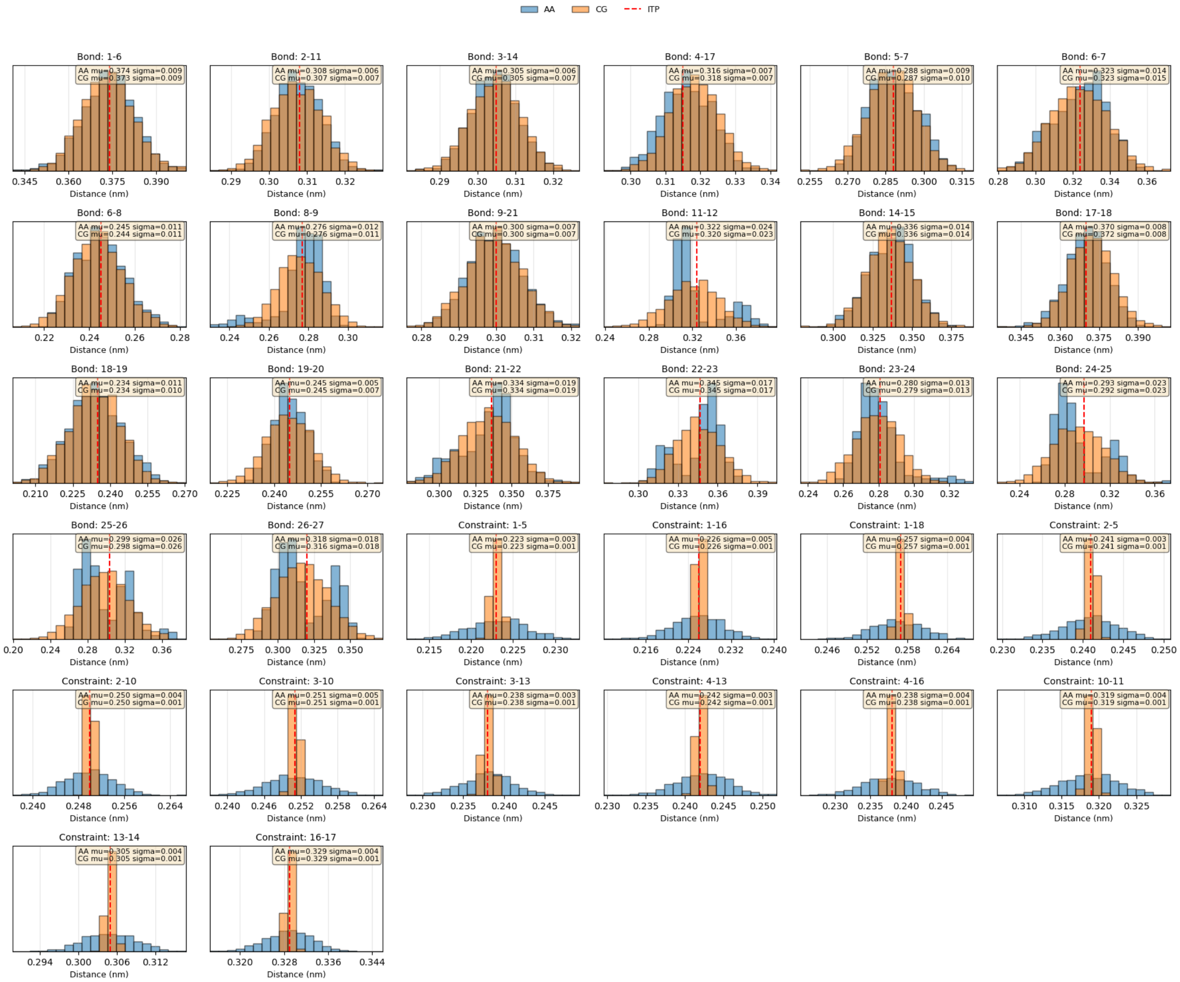
AA
CG
ITP
Bond: 1-6
AA mu=0.374 sigma=0.009
CG mu=0.373 sigma=0.009
Bond: 2-11
AA mu=0.308 sigma=0.006
CG mu=0.307 sigma=0.007
Bond: 3-14
AA mu=0.305 sigma=0.006
CG mu=0.305 sigma=0.007
Bond: 4-17
AA mu=0.316 sigma=0.007
CG mu=0.318 sigma=0.007
Bond: 5-7
AA mu=0.288 sigma=0.009
CG mu=0.287 sigma=0.010
Bond: 6-7
AA mu=0.323 sigma=0.014
CG mu=0.323 sigma=0.015
Bond: 6-8
AA mu=0.245 sigma=0.011
CG mu=0.244 sigma=0.011
Bond: 8-9
AA mu=0.276 sigma=0.012
CG mu=0.276 sigma=0.011
Bond: 9-21
AA mu=0.300 sigma=0.007
CG mu=0.300 sigma=0.007
Bond: 11-12
AA mu=0.322 sigma=0.024
CG mu=0.320 sigma=0.023
Bond: 14-15
AA mu=0.336 sigma=0.014
CG mu=0.336 sigma=0.014
Bond: 17-18
AA mu=0.370 sigma=0.008
CG mu=0.372 sigma=0.008
Bond: 18-19
AA mu=0.234 sigma=0.011
CG mu=0.234 sigma=0.010
Bond: 19-20
AA mu=0.245 sigma=0.005
CG mu=0.245 sigma=0.007
Bond: 21-22
AA mu=0.334 sigma=0.019
CG mu=0.334 sigma=0.019
Bond: 22-23
AA mu=0.345 sigma=0.017
CG mu=0.345 sigma=0.017
Bond: 23-24
AA mu=0.280 sigma=0.013
CG mu=0.279 sigma=0.013
Bond: 24-25
AA mu=0.293 sigma=0.023
CG mu=0.292 sigma=0.023
Bond: 25-26
AA mu=0.299 sigma=0.026
CG mu=0.298 sigma=0.026
Bond: 26-27
AA mu=0.318 sigma=0.018
CG mu=0.316 sigma=0.018
Constraint: 1-5
AA mu=0.223 sigma=0.003
CG mu=0.223 sigma=0.001
Constraint: 1-16
AA mu=0.226 sigma=0.005
CG mu=0.226 sigma=0.001
Constraint: 1-18
AA mu=0.257 sigma=0.004
CG mu=0.257 sigma=0.001
Constraint: 2-5
AA mu=0.241 sigma=0.003
CG mu=0.241 sigma=0.001
Constraint: 2-10
AA mu=0.250 sigma=0.004
CG mu=0.250 sigma=0.001
Constraint: 3-10
AA mu=0.251 sigma=0.005
CG mu=0.251 sigma=0.001
Constraint: 3-13
AA mu=0.238 sigma=0.003
CG mu=0.238 sigma=0.001
Constraint: 4-13
AA mu=0.242 sigma=0.003
CG mu=0.242 sigma=0.001
Constraint: 4-16
AA mu=0.238 sigma=0.004
CG mu=0.238 sigma=0.001
Constraint: 10-11
AA mu=0.319 sigma=0.004
CG mu=0.319 sigma=0.001
Constraint: 13-14
AA mu=0.305 sigma=0.004
CG mu=0.305 sigma=0.001
Constraint: 16-17
AA mu=0.329 sigma=0.004
CG mu=0.329 sigma=0.001
Distance (nm)

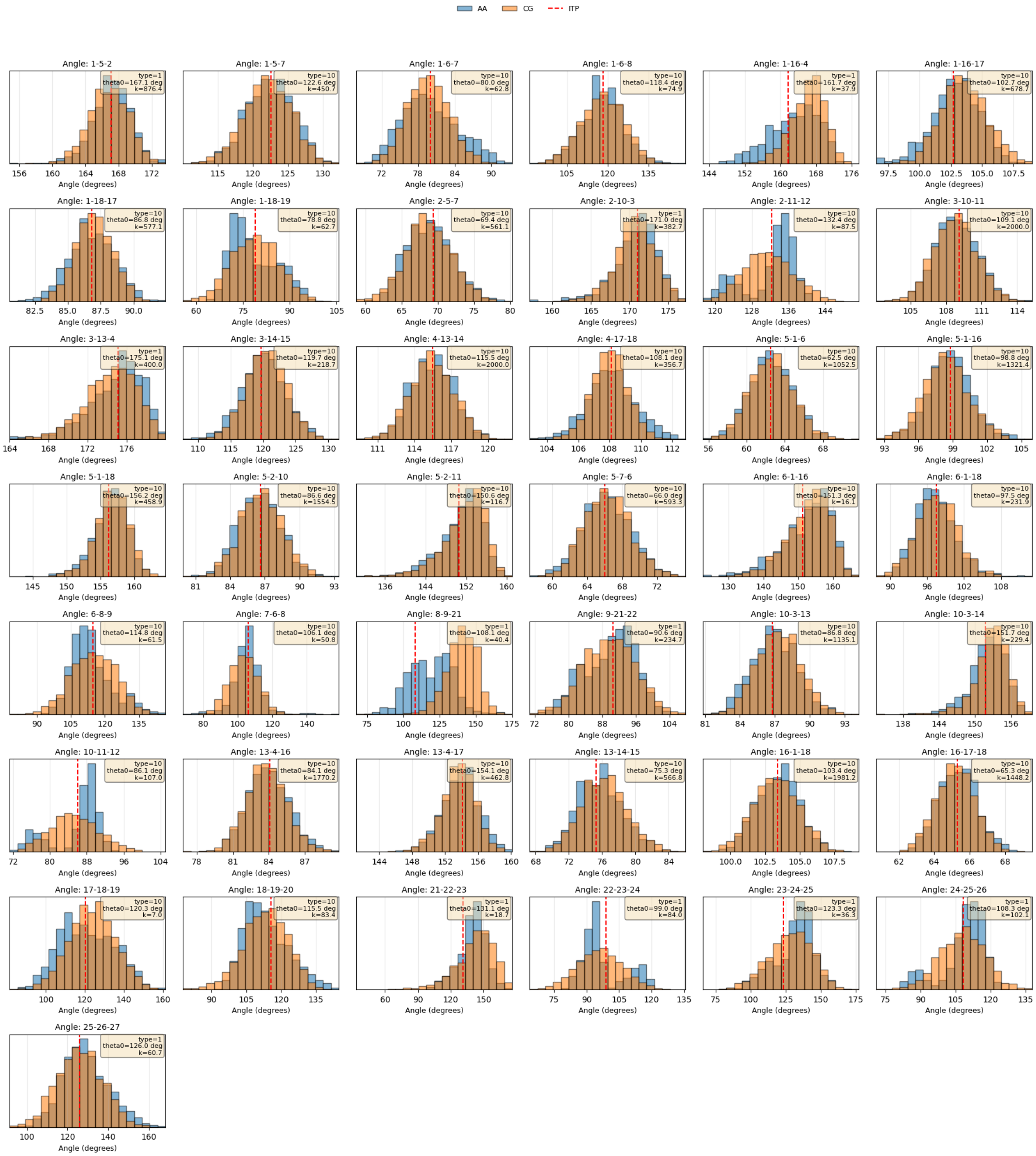
AA
CG
ITP
Angle: 1-5-2
type=1 theta0=167.1 deg k=876.4
Angle: 1-5-7
type=10 theta0=122.6 deg k=450.7
Angle: 1-6-7
type=10 theta0=80.0 deg k=62.8
Angle: 1-6-8
type=10 theta0=118.4 deg k=74.9
Angle: 1-16-4
type=1 theta0=161.7 deg k=37.9
Angle: 1-16-17
type=10 theta0=102.7 deg k=678.7
Angle: 1-18-17
type=10 theta0=86.8 deg k=577.1
Angle: 1-18-19
type=10 theta0=78.8 deg k=62.7
Angle: 2-5-7
type=10 theta0=69.4 deg k=561.1
Angle: 2-10-3
type=1 theta0=171.0 deg k=382.7
Angle: 2-11-12
type=10 theta0=132.4 deg k=87.5
Angle: 3-10-11
type=10 theta0=109.1 deg k=2000.0
Angle: 3-13-4
type=1 theta0=175.1 deg k=400.0
Angle: 3-14-15
type=10 theta0=119.7 deg k=218.7
Angle: 4-13-14
type=10 theta0=115.5 deg k=2000.0
Angle: 4-17-18
type=10 theta0=108.1 deg k=356.7
Angle: 5-1-6
type=10 theta0=62.5 deg k=1052.5
Angle: 5-1-16
type=10 theta0=98.8 deg k=1321.4
Angle: 5-1-18
type=10 theta0=156.2 deg k=458.9
Angle: 5-2-10
type=10 theta0=86.6 deg k=1554.5
Angle: 5-2-11
type=10 theta0=150.6 deg k=116.7
Angle: 5-7-6
type=10 theta0=66.0 deg k=593.3
Angle: 6-1-16
type=10 theta0=151.3 deg k=16.1
Angle: 6-1-18
type=10 theta0=97.5 deg k=231.9
Angle: 6-8-9
type=10 theta0=114.8 deg k=61.5
Angle: 7-6-8
type=10 theta0=106.1 deg k=50.8
Angle: 8-9-21
type=1 theta0=108.1 deg k=40.4
Angle: 9-21-22
type=1 theta0=90.6 deg k=234.7
Angle: 10-3-13
type=10 theta0=86.8 deg k=1135.1
Angle: 10-3-14
type=10 theta0=151.7 deg k=229.4
Angle: 10-11-12
type=10 theta0=86.1 deg k=107.0
Angle: 13-4-16
type=10 theta0=84.1 deg k=1770.2
Angle: 13-4-17
type=10 theta0=154.1 deg k=462.8
Angle: 13-14-15
type=10 theta0=75.3 deg k=566.8
Angle: 16-1-18
type=10 theta0=103.4 deg k=1981.2
Angle: 16-17-18
type=10 theta0=65.3 deg k=1448.2
Angle: 17-18-19
type=10 theta0=120.3 deg k=7.0
Angle: 18-19-20
type=10 theta0=115.5 deg k=83.4
Angle: 21-22-23
type=1 theta0=131.1 deg k=18.7
Angle: 22-23-24
type=1 theta0=99.0 deg k=84.0
Angle: 23-24-25
type=1 theta0=123.3 deg k=36.3
Angle: 24-25-26
type=1 theta0=108.3 deg k=102.1
Angle: 25-26-27
type=1 theta0=126.0 deg k=60.7
Angle (degrees)

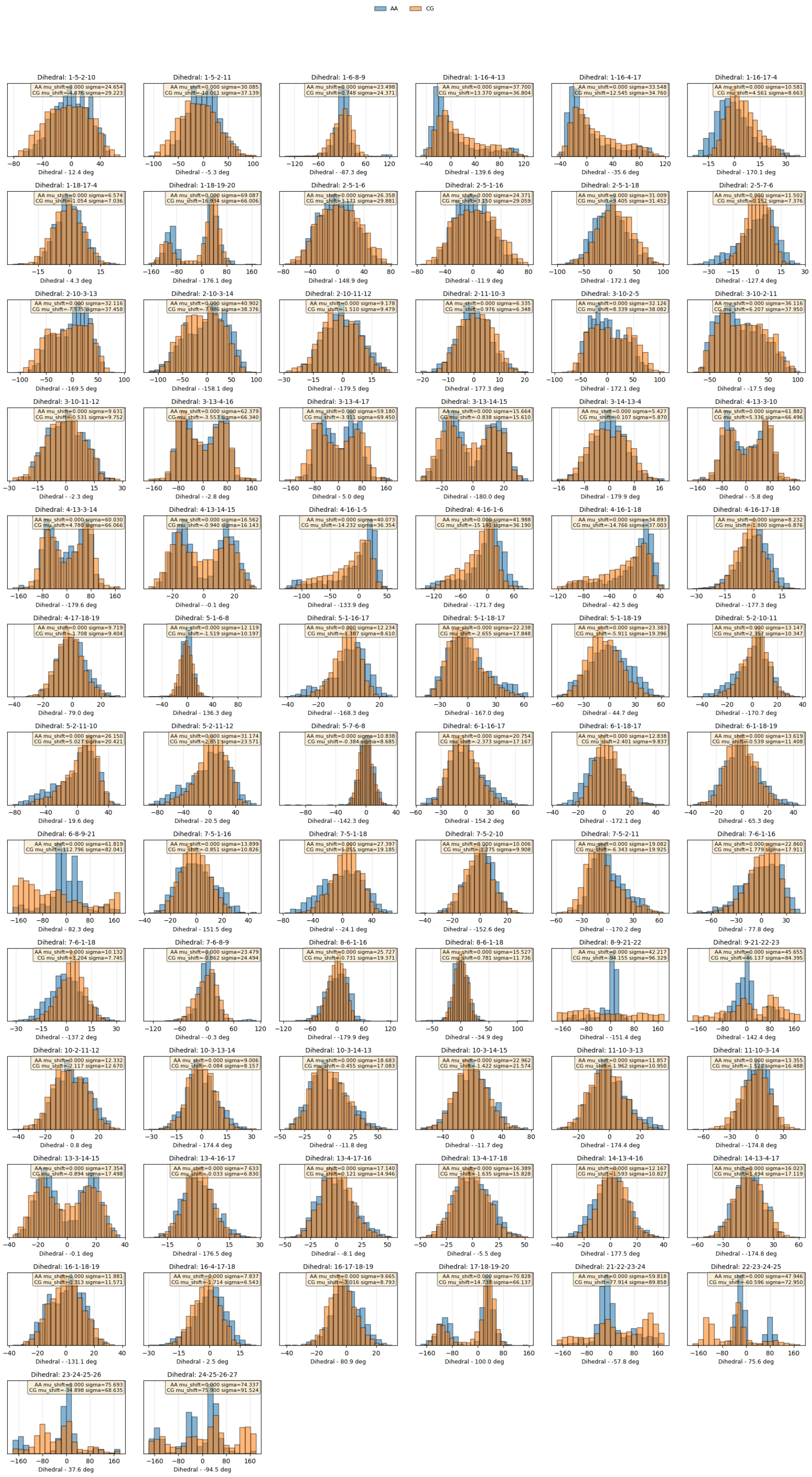
AA
CG
Dihedral: 1-5-2-10
Dihedral - 12.4 deg
Dihedral: 1-5-2-11
Dihedral - -5.3 deg
Dihedral: 1-6-8-9
Dihedral - -87.3 deg
Dihedral: 1-16-4-13
Dihedral - 139.6 deg
Dihedral: 1-16-4-17
Dihedral - -35.6 deg
Dihedral: 1-16-17-4
Dihedral - 170.1 deg
Dihedral: 1-18-17-4
Dihedral - 4.3 deg
Dihedral: 1-18-19-20
Dihedral - 176.1 deg
Dihedral: 2-5-1-6
Dihedral - 148.9 deg
Dihedral: 2-5-1-16
Dihedral - -11.9 deg
Dihedral: 2-5-1-18
Dihedral - 172.1 deg
Dihedral: 2-5-7-6
Dihedral - -127.4 deg
Dihedral: 2-10-3-13
Dihedral - -169.5 deg
Dihedral: 2-10-3-14
Dihedral - -158.1 deg
Dihedral: 2-10-11-12
Dihedral - -179.5 deg
Dihedral: 2-11-10-3
Dihedral - 177.3 deg
Dihedral: 3-10-2-5
Dihedral - 172.1 deg
Dihedral: 3-10-2-11
Dihedral - -17.5 deg
Dihedral: 3-10-11-12
Dihedral - -2.3 deg
Dihedral: 3-13-4-16
Dihedral - -2.8 deg
Dihedral: 3-13-4-17
Dihedral - 5.0 deg
Dihedral: 3-13-14-15
Dihedral - -180.0 deg
Dihedral: 3-14-13-4
Dihedral - 179.9 deg
Dihedral: 4-13-3-10
Dihedral - -5.8 deg
Dihedral: 4-13-3-14
Dihedral - -179.6 deg
Dihedral: 4-13-14-15
Dihedral - -0.1 deg
Dihedral: 4-16-1-5
Dihedral - -133.9 deg
Dihedral: 4-16-1-6
Dihedral - -171.7 deg
Dihedral: 4-16-1-18
Dihedral - 42.5 deg
Dihedral: 4-16-17-18
Dihedral - -177.3 deg
Dihedral: 4-17-18-19
Dihedral - 79.0 deg
Dihedral: 5-1-6-8
Dihedral - 136.3 deg
Dihedral: 5-1-16-17
Dihedral - -168.3 deg
Dihedral: 5-1-18-17
Dihedral - 167.0 deg
Dihedral: 5-1-18-19
Dihedral - 44.7 deg
Dihedral: 5-2-10-11
Dihedral - -170.7 deg
Dihedral: 5-2-11-10
Dihedral - 19.6 deg
Dihedral: 5-2-11-12
Dihedral - 20.5 deg
Dihedral: 5-7-6-8
Dihedral - -142.3 deg
Dihedral: 6-1-16-17
Dihedral - 154.2 deg
Dihedral: 6-1-18-17
Dihedral - -172.1 deg
Dihedral: 6-1-18-19
Dihedral - 65.3 deg
Dihedral: 6-8-9-21
Dihedral - 82.3 deg
Dihedral: 7-5-1-16
Dihedral - 151.5 deg
Dihedral: 7-5-1-18
Dihedral - -24.1 deg
Dihedral: 7-5-2-10
Dihedral - -152.6 deg
Dihedral: 7-5-2-11
Dihedral - -170.2 deg
Dihedral: 7-6-1-16
Dihedral - 77.8 deg
Dihedral: 7-6-1-18
Dihedral - -137.2 deg
Dihedral: 7-6-8-9
Dihedral - -0.3 deg
Dihedral: 8-6-1-16
Dihedral - -179.9 deg
Dihedral: 8-6-1-18
Dihedral - -34.9 deg
Dihedral: 8-9-21-22
Dihedral - -151.4 deg
Dihedral: 9-21-22-23
Dihedral - 142.4 deg
Dihedral: 10-2-11-12
Dihedral - 0.8 deg
Dihedral: 10-3-13-14
Dihedral - 174.4 deg
Dihedral: 10-3-14-13
Dihedral - -11.8 deg
Dihedral: 10-3-14-15
Dihedral - -11.7 deg
Dihedral: 11-10-3-13
Dihedral - 174.4 deg
Dihedral: 11-10-3-14
Dihedral - -174.8 deg
Dihedral: 13-3-14-15
Dihedral - -0.1 deg
Dihedral: 13-4-16-17
Dihedral - 176.5 deg
Dihedral: 13-4-17-16
Dihedral - -8.1 deg
Dihedral: 13-4-17-18
Dihedral - -5.5 deg
Dihedral: 14-13-4-16
Dihedral - 177.5 deg
Dihedral: 14-13-4-17
Dihedral - -174.8 deg
Dihedral: 16-1-18-19
Dihedral - -131.1 deg
Dihedral: 16-4-17-18
Dihedral - 2.5 deg
Dihedral: 16-17-18-19
Dihedral - 80.9 deg
Dihedral: 17-18-19-20
Dihedral - 100.0 deg
Dihedral: 21-22-23-24
Dihedral - -57.8 deg
Dihedral: 22-23-24-25
Dihedral - 75.6 deg
Dihedral: 23-24-25-26
Dihedral - 37.6 deg
Dihedral: 24-25-26-27
Dihedral - -94.5 deg

### DMBI

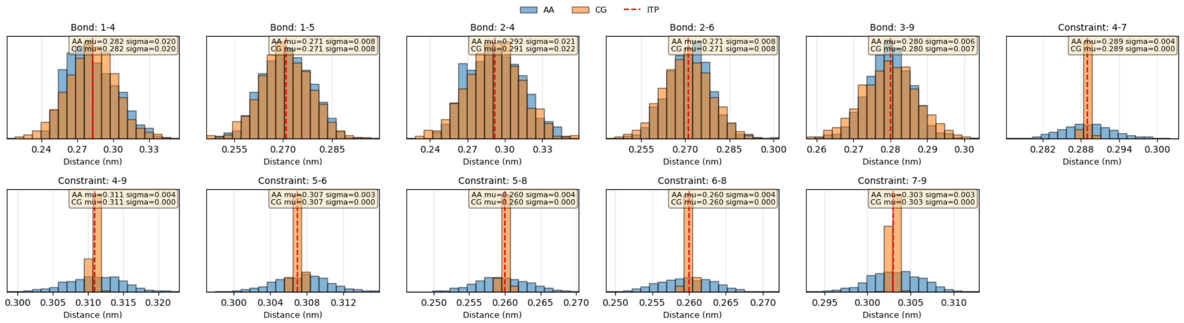

AA
CG
ITP
Bond: 1-4
AA mu=0.282 sigma=0.020
CG mu=0.282 sigma=0.020
Bond: 1-5
AA mu=0.271 sigma=0.008
CG mu=0.271 sigma=0.008
Bond: 2-4
AA mu=0.292 sigma=0.021
CG mu=0.291 sigma=0.022
Bond: 2-6
AA mu=0.271 sigma=0.008
CG mu=0.271 sigma=0.008
Bond: 3-9
AA mu=0.280 sigma=0.006
CG mu=0.280 sigma=0.007
Constraint: 4-7
AA mu=0.289 sigma=0.004
CG mu=0.289 sigma=0.000
Constraint: 4-9
AA mu=0.311 sigma=0.004
CG mu=0.311 sigma=0.000
Constraint: 5-6
AA mu=0.307 sigma=0.003
CG mu=0.307 sigma=0.000
Constraint: 5-8
AA mu=0.260 sigma=0.004
CG mu=0.260 sigma=0.000
Constraint: 6-8
AA mu=0.260 sigma=0.004
CG mu=0.260 sigma=0.000
Constraint: 7-9
AA mu=0.303 sigma=0.003
CG mu=0.303 sigma=0.000
Distance (nm)


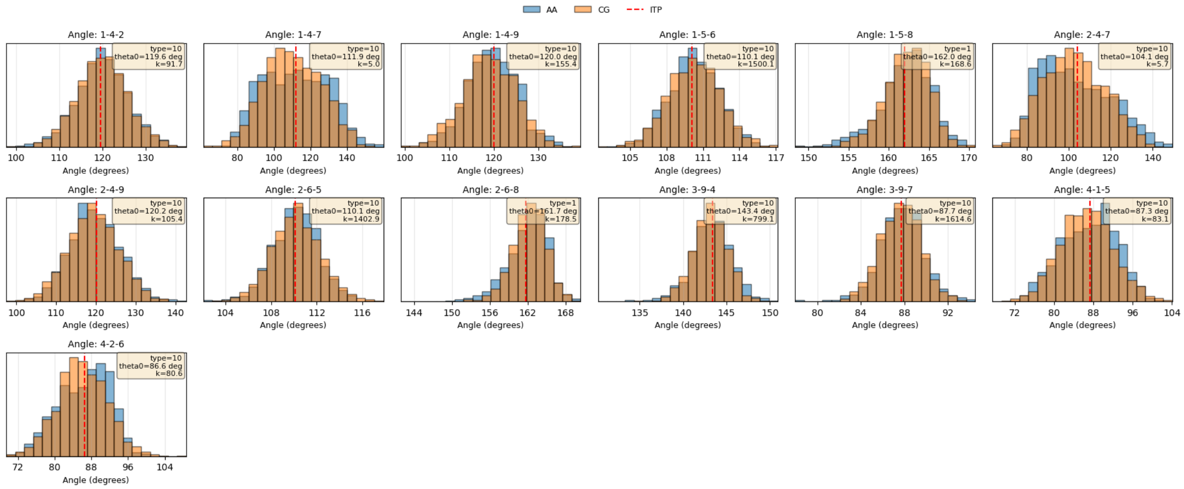

AA
CG
ITP
Angle: 1-4-2
type=10
theta0=119.6 deg
k=91.7
Angle: 1-4-7
type=10
theta0=111.9 deg
k=5.0
Angle: 1-4-9
type=10
theta0=120.0 deg
k=155.4
Angle: 1-5-6
type=10
theta0=110.1 deg
k=1500.1
Angle: 1-5-8
type=1
theta0=162.0 deg
k=168.6
Angle: 2-4-7
type=10
theta0=104.1 deg
k=5.7
Angle: 2-4-9
type=10
theta0=120.2 deg
k=105.4
Angle: 2-6-5
type=10
theta0=110.1 deg
k=1402.9
Angle: 2-6-8
type=1
theta0=161.7 deg
k=178.5
Angle: 3-9-4
type=10
theta0=143.4 deg
k=799.1
Angle: 3-9-7
type=10
theta0=87.7 deg
k=1614.6
Angle: 4-1-5
type=10
theta0=87.3 deg
k=83.1
Angle: 4-2-6
type=10
theta0=86.6 deg
k=80.6
Angle (degrees)

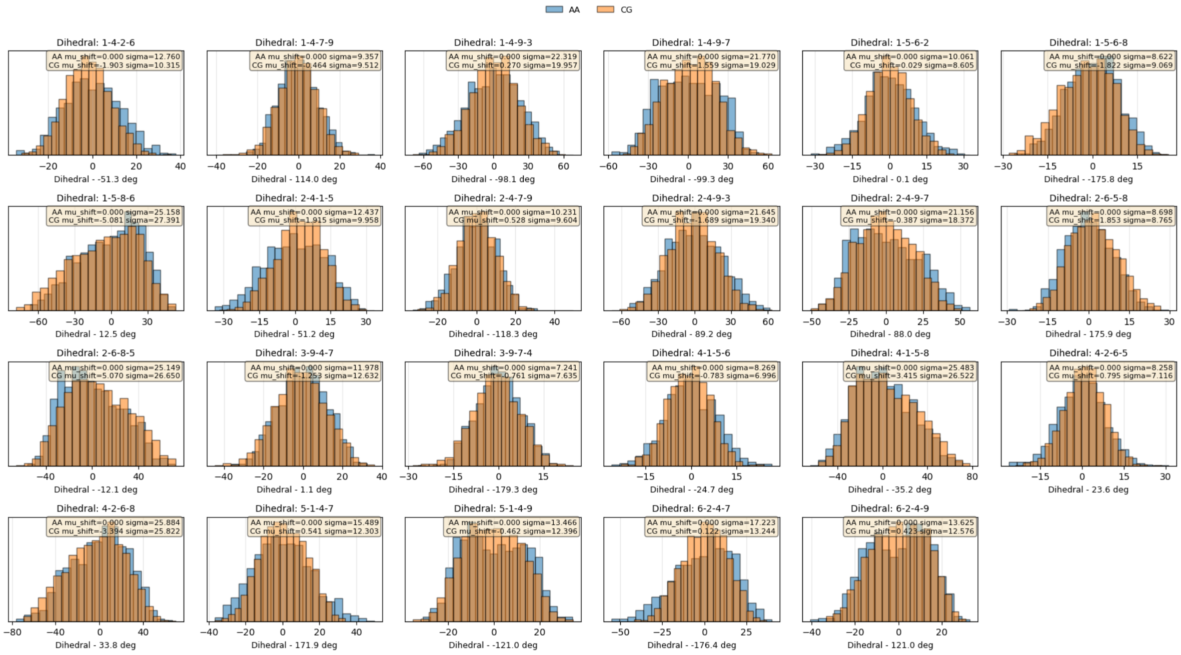
AA
CG
Dihedral: 1-4-2-6
AA mu_shift=0.000 sigma=12.760
CG mu_shift=-1.903 sigma=10.315
Dihedral - -51.3 deg
Dihedral: 1-4-7-9
AA mu_shift=0.000 sigma=9.357
CG mu_shift=0.464 sigma=9.512
Dihedral - 114.0 deg
Dihedral: 1-4-9-3
AA mu_shift=0.000 sigma=22.319
CG mu_shift=0.270 sigma=19.957
Dihedral - -98.1 deg
Dihedral: 1-4-9-7
AA mu_shift=0.000 sigma=21.770
CG mu_shift=-1.559 sigma=19.029
Dihedral - -99.3 deg
Dihedral: 1-5-6-2
AA mu_shift=0.000 sigma=10.061
CG mu_shift=0.029 sigma=8.605
Dihedral - 0.1 deg
Dihedral: 1-5-6-8
AA mu_shift=0.000 sigma=8.622
CG mu_shift=-1.822 sigma=9.069
Dihedral - -175.8 deg
Dihedral: 1-5-8-6
AA mu_shift=0.000 sigma=25.158
CG mu_shift=-5.081 sigma=27.391
Dihedral - 12.5 deg
Dihedral: 2-4-1-5
AA mu_shift=0.000 sigma=12.437
CG mu_shift=1.915 sigma=9.958
Dihedral - 51.2 deg
Dihedral: 2-4-7-9
AA mu_shift=0.000 sigma=10.231
CG mu_shift=0.528 sigma=9.604
Dihedral - -118.3 deg
Dihedral: 2-4-9-3
AA mu_shift=0.000 sigma=21.645
CG mu_shift=-1.689 sigma=19.340
Dihedral - 89.2 deg
Dihedral: 2-4-9-7
AA mu_shift=0.000 sigma=21.156
CG mu_shift=-0.387 sigma=18.372
Dihedral - 88.0 deg
Dihedral: 2-6-5-8
AA mu_shift=0.000 sigma=8.698
CG mu_shift=1.853 sigma=8.765
Dihedral - 175.9 deg
Dihedral: 2-6-8-5
AA mu_shift=0.000 sigma=25.149
CG mu_shift=5.070 sigma=26.650
Dihedral - -12.1 deg
Dihedral: 3-9-4-7
AA mu_shift=0.000 sigma=11.978
CG mu_shift=-1.253 sigma=12.632
Dihedral - 1.1 deg
Dihedral: 3-9-7-4
AA mu_shift=0.000 sigma=7.241
CG mu_shift=-0.761 sigma=7.635
Dihedral - -179.3 deg
Dihedral: 4-1-5-6
AA mu_shift=0.000 sigma=8.269
CG mu_shift=-0.783 sigma=6.996
Dihedral - -24.7 deg
Dihedral: 4-1-5-8
AA mu_shift=0.000 sigma=25.483
CG mu_shift=3.415 sigma=26.522
Dihedral - -35.2 deg
Dihedral: 4-2-6-5
AA mu_shift=0.000 sigma=8.258
CG mu_shift=0.795 sigma=7.116
Dihedral - 23.6 deg
Dihedral: 4-2-6-8
AA mu_shift=0.000 sigma=25.884
CG mu_shift=-3.394 sigma=25.822
Dihedral - 33.8 deg
Dihedral: 5-1-4-7
AA mu_shift=0.000 sigma=15.489
CG mu_shift=0.541 sigma=12.303
Dihedral - 171.9 deg
Dihedral: 5-1-4-9
AA mu_shift=0.000 sigma=13.466
CG mu_shift=-0.462 sigma=12.396
Dihedral - -121.0 deg
Dihedral: 6-2-4-7
AA mu_shift=0.000 sigma=17.223
CG mu_shift=0.122 sigma=13.244
Dihedral - -176.4 deg
Dihedral: 6-2-4-9
AA mu_shift=0.000 sigma=13.625
CG mu_shift=0.423 sigma=12.576
Dihedral - 121.0 deg

### FTA

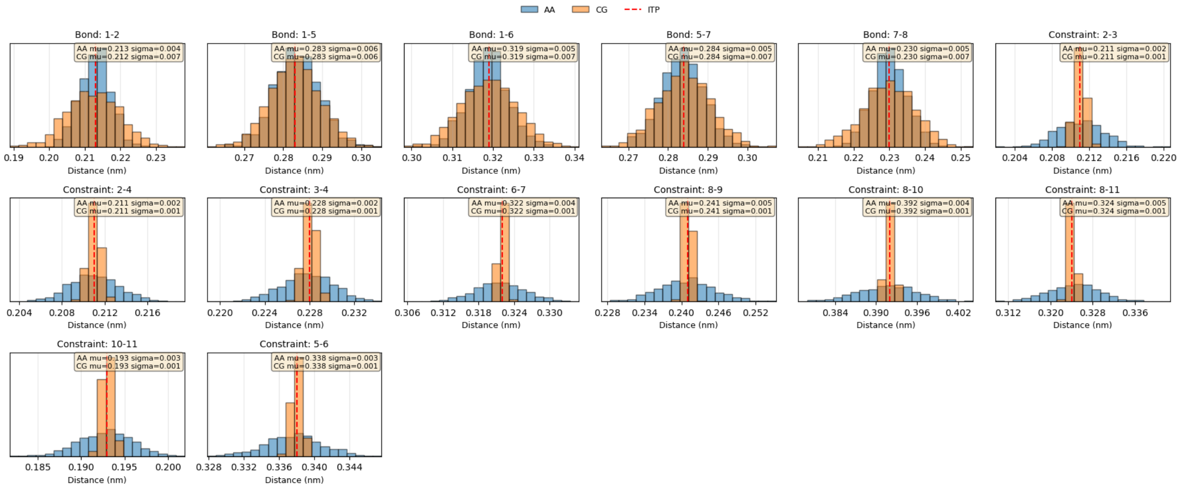
AA
CG
ITP
Bond: 1-2
AA mu=0.213 sigma=0.004
CG mu=0.212 sigma=0.007
Bond: 1-5
AA mu=0.283 sigma=0.006
CG mu=0.283 sigma=0.006
Bond: 1-6
AA mu=0.319 sigma=0.005
CG mu=0.319 sigma=0.007
Bond: 5-7
AA mu=0.284 sigma=0.005
CG mu=0.284 sigma=0.007
Bond: 7-8
AA mu=0.230 sigma=0.005
CG mu=0.230 sigma=0.007
Constraint: 2-3
AA mu=0.211 sigma=0.002
CG mu=0.211 sigma=0.001
Constraint: 2-4
AA mu=0.211 sigma=0.002
CG mu=0.211 sigma=0.001
Constraint: 3-4
AA mu=0.228 sigma=0.002
CG mu=0.228 sigma=0.001
Constraint: 6-7
AA mu=0.322 sigma=0.004
CG mu=0.322 sigma=0.001
Constraint: 8-9
AA mu=0.241 sigma=0.005
CG mu=0.241 sigma=0.001
Constraint: 8-10
AA mu=0.392 sigma=0.004
CG mu=0.392 sigma=0.001
Constraint: 8-11
AA mu=0.324 sigma=0.005
CG mu=0.324 sigma=0.001
Constraint: 10-11
AA mu=0.193 sigma=0.003
CG mu=0.193 sigma=0.001
Constraint: 5-6
AA mu=0.338 sigma=0.003
CG mu=0.338 sigma=0.001
Distance (nm)

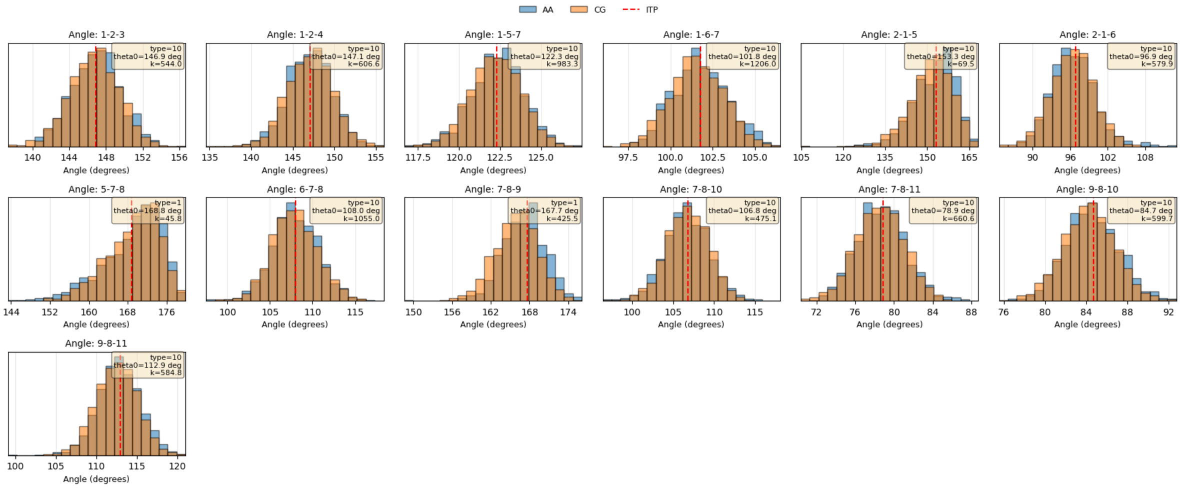
AA
CG
ITP
Angle: 1-2-3
Angle: 1-2-4
Angle: 1-5-7
Angle: 1-6-7
Angle: 2-1-5
Angle: 2-1-6
Angle: 5-7-8
Angle: 6-7-8
Angle: 7-8-9
Angle: 7-8-10
Angle: 7-8-11
Angle: 9-8-10
Angle: 9-8-11
Angle (degrees)

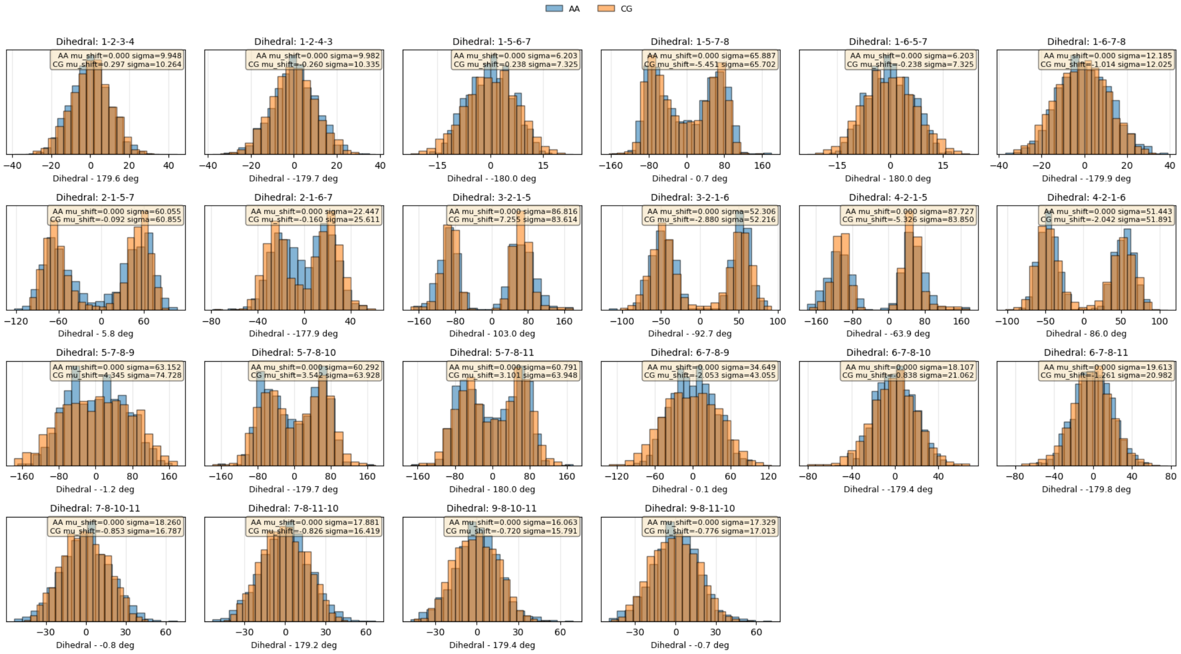
AA
CG
Dihedral: 1-2-3-4
Dihedral: 1-2-4-3
Dihedral: 1-5-6-7
Dihedral: 1-5-7-8
Dihedral: 1-6-5-7
Dihedral: 1-6-7-8
Dihedral: 2-1-5-7
Dihedral: 2-1-6-7
Dihedral: 3-2-1-5
Dihedral: 3-2-1-6
Dihedral: 4-2-1-5
Dihedral: 4-2-1-6
Dihedral: 5-7-8-9
Dihedral: 5-7-8-10
Dihedral: 5-7-8-11
Dihedral: 6-7-8-9
Dihedral: 6-7-8-10
Dihedral: 6-7-8-11
Dihedral: 7-8-10-11
Dihedral: 7-8-11-10
Dihedral: 9-8-10-11
Dihedral: 9-8-11-10

### HEM

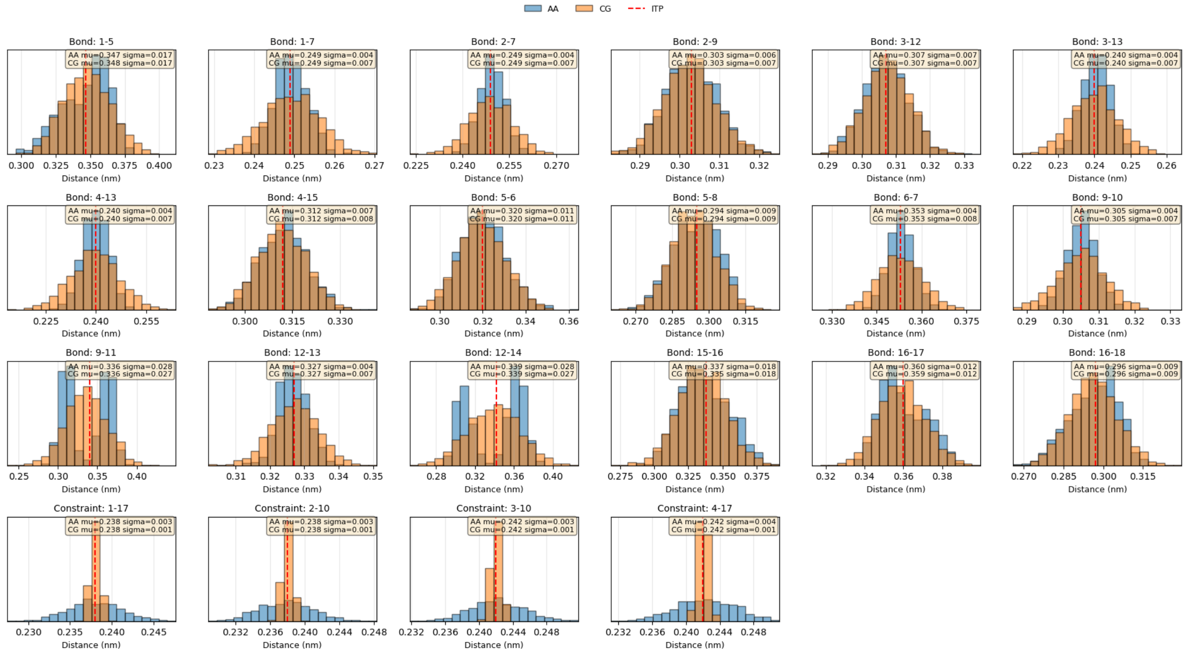
AA
CG
ITP
Bond: 1-5
AA mu=0.347 sigma=0.017
CG mu=0.348 sigma=0.017
Bond: 1-7
AA mu=0.249 sigma=0.004
CG mu=0.249 sigma=0.007
Bond: 2-7
AA mu=0.249 sigma=0.004
CG mu=0.249 sigma=0.007
Bond: 2-9
AA mu=0.303 sigma=0.006
CG mu=0.303 sigma=0.007
Bond: 3-12
AA mu=0.307 sigma=0.007
CG mu=0.307 sigma=0.007
Bond: 3-13
AA mu=0.240 sigma=0.004
CG mu=0.240 sigma=0.007
Bond: 4-13
AA mu=0.240 sigma=0.004
CG mu=0.240 sigma=0.007
Bond: 4-15
AA mu=0.312 sigma=0.007
CG mu=0.312 sigma=0.008
Bond: 5-6
AA mu=0.320 sigma=0.011
CG mu=0.320 sigma=0.011
Bond: 5-8
AA mu=0.294 sigma=0.009
CG mu=0.294 sigma=0.009
Bond: 6-7
AA mu=0.353 sigma=0.004
CG mu=0.353 sigma=0.008
Bond: 9-10
AA mu=0.305 sigma=0.004
CG mu=0.305 sigma=0.007
Bond: 9-11
AA mu=0.336 sigma=0.028
CG mu=0.336 sigma=0.027
Bond: 12-13
AA mu=0.327 sigma=0.004
CG mu=0.327 sigma=0.007
Bond: 12-14
AA mu=0.339 sigma=0.028
CG mu=0.339 sigma=0.027
Bond: 15-16
AA mu=0.337 sigma=0.018
CG mu=0.335 sigma=0.018
Bond: 16-17
AA mu=0.360 sigma=0.012
CG mu=0.359 sigma=0.012
Bond: 16-18
AA mu=0.296 sigma=0.009
CG mu=0.296 sigma=0.009
Constraint: 1-17
AA mu=0.238 sigma=0.003
CG mu=0.238 sigma=0.001
Constraint: 2-10
AA mu=0.238 sigma=0.003
CG mu=0.238 sigma=0.001
Constraint: 3-10
AA mu=0.242 sigma=0.003
CG mu=0.242 sigma=0.001
Constraint: 4-17
AA mu=0.242 sigma=0.004
CG mu=0.242 sigma=0.001
Distance (nm)

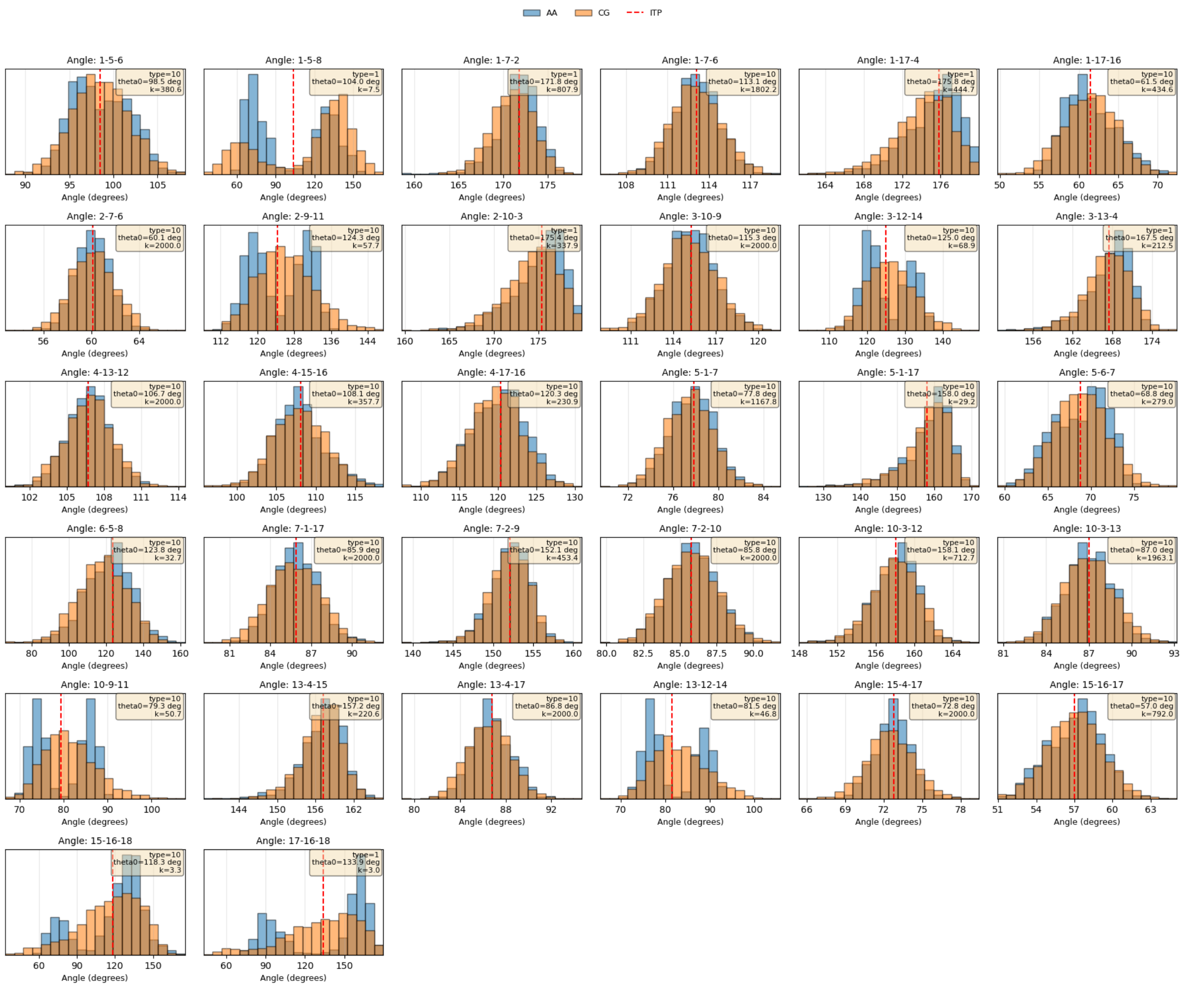
AA
CG
ITP
Angle: 1-5-6
type=10 theta0=98.5 deg k=380.6
Angle: 1-5-8
type=1 theta0=104.0 deg k=7.5
Angle: 1-7-2
type=1 theta0=171.8 deg k=807.9
Angle: 1-7-6
type=10 theta0=113.1 deg k=1802.2
Angle: 1-17-4
type=1 theta0=175.8 deg k=444.7
Angle: 1-17-16
type=10 theta0=61.5 deg k=434.6
Angle: 2-7-6
type=10 theta0=60.1 deg k=2000.0
Angle: 2-9-11
type=10 theta0=124.3 deg k=57.7
Angle: 2-10-3
type=1 theta0=175.4 deg k=337.9
Angle: 3-10-9
type=10 theta0=115.3 deg k=2000.0
Angle: 3-12-14
type=10 theta0=125.0 deg k=68.9
Angle: 3-13-4
type=1 theta0=167.5 deg k=212.5
Angle: 4-13-12
type=10 theta0=106.7 deg k=2000.0
Angle: 4-15-16
type=10 theta0=108.1 deg k=357.7
Angle: 4-17-16
type=10 theta0=120.3 deg k=230.9
Angle: 5-1-7
type=10 theta0=77.8 deg k=1167.8
Angle: 5-1-17
type=10 theta0=158.0 deg k=29.2
Angle: 5-6-7
type=10 theta0=68.8 deg k=279.0
Angle: 6-5-8
type=10 theta0=123.8 deg k=32.7
Angle: 7-1-17
type=10 theta0=85.9 deg k=2000.0
Angle: 7-2-9
type=10 theta0=152.1 deg k=453.4
Angle: 7-2-10
type=10 theta0=85.8 deg k=2000.0
Angle: 10-3-12
type=10 theta0=158.1 deg k=712.7
Angle: 10-3-13
type=10 theta0=87.0 deg k=1963.1
Angle: 10-9-11
type=10 theta0=79.3 deg k=50.7
Angle: 13-4-15
type=10 theta0=157.2 deg k=220.6
Angle: 13-4-17
type=10 theta0=86.8 deg k=2000.0
Angle: 13-12-14
type=10 theta0=81.5 deg k=46.8
Angle: 15-4-17
type=10 theta0=72.8 deg k=2000.0
Angle: 15-16-17
type=10 theta0=57.0 deg k=792.0
Angle: 15-16-18
type=10 theta0=118.3 deg k=3.3
Angle: 17-16-18
type=1 theta0=133.9 deg k=3.0
Angle (degrees)

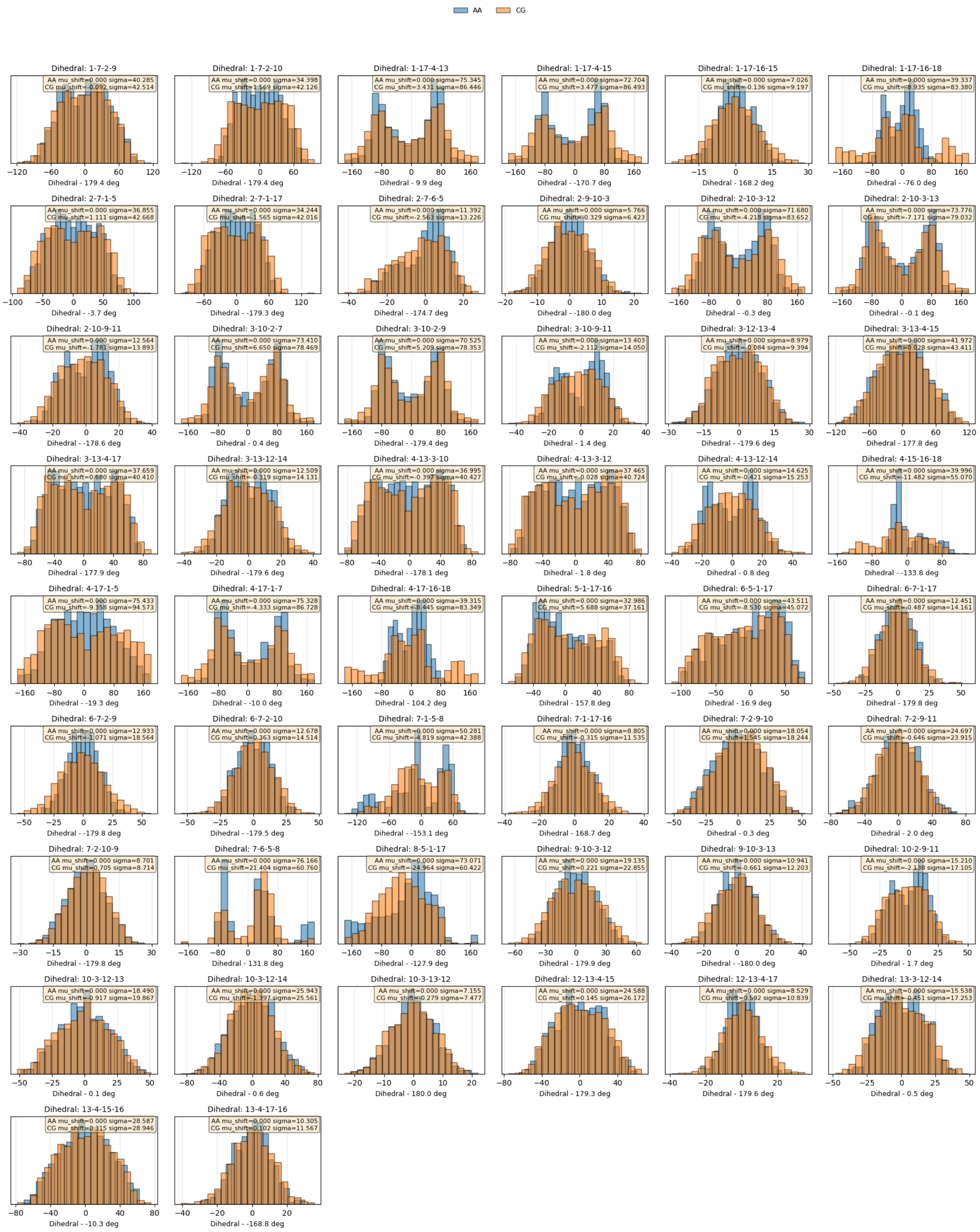
AA
CG
Dihedral: 1-7-2-9
AA mu_shift=0.000 sigma=40.285
CG mu_shift=-0.092 sigma=42.514
Dihedral - 179.4 deg
Dihedral: 1-7-2-10
AA mu_shift=0.000 sigma=34.398
CG mu_shift=1.569 sigma=42.126
Dihedral - 179.4 deg
Dihedral: 1-17-4-13
AA mu_shift=0.000 sigma=75.345
CG mu_shift=3.431 sigma=86.446
Dihedral - 9.9 deg
Dihedral: 1-17-4-15
AA mu_shift=0.000 sigma=72.704
CG mu_shift=3.477 sigma=86.493
Dihedral - -170.7 deg
Dihedral: 1-17-16-15
AA mu_shift=0.000 sigma=7.026
CG mu_shift=-0.136 sigma=9.197
Dihedral - 168.2 deg
Dihedral: 1-17-16-18
AA mu_shift=0.000 sigma=39.337
CG mu_shift=-8.935 sigma=83.380
Dihedral - -76.0 deg
Dihedral: 2-7-1-5
AA mu_shift=0.000 sigma=36.855
CG mu_shift=1.111 sigma=42.668
Dihedral - -3.7 deg
Dihedral: 2-7-1-17
AA mu_shift=0.000 sigma=34.244
CG mu_shift=-1.565 sigma=42.016
Dihedral - -179.3 deg
Dihedral: 2-7-6-5
AA mu_shift=0.000 sigma=11.392
CG mu_shift=-2.563 sigma=13.226
Dihedral - -174.7 deg
Dihedral: 2-9-10-3
AA mu_shift=0.000 sigma=5.766
CG mu_shift=-0.329 sigma=6.423
Dihedral - -180.0 deg
Dihedral: 2-10-3-12
AA mu_shift=0.000 sigma=71.680
CG mu_shift=-4.218 sigma=83.652
Dihedral - -0.3 deg
Dihedral: 2-10-3-13
AA mu_shift=0.000 sigma=73.776
CG mu_shift=-7.171 sigma=79.032
Dihedral - -0.1 deg
Dihedral: 2-10-9-11
AA mu_shift=0.000 sigma=12.564
CG mu_shift=-1.781 sigma=13.893
Dihedral - -178.6 deg
Dihedral: 3-10-2-7
AA mu_shift=0.000 sigma=73.410
CG mu_shift=6.650 sigma=78.469
Dihedral - 0.4 deg
Dihedral: 3-10-2-9
AA mu_shift=0.000 sigma=70.525
CG mu_shift=5.209 sigma=78.353
Dihedral - -179.4 deg
Dihedral: 3-10-9-11
AA mu_shift=0.000 sigma=13.403
CG mu_shift=-2.112 sigma=14.050
Dihedral - 1.4 deg
Dihedral: 3-12-13-4
AA mu_shift=0.000 sigma=8.979
CG mu_shift=-0.084 sigma=9.394
Dihedral - -179.6 deg
Dihedral: 3-13-4-15
AA mu_shift=0.000 sigma=41.972
CG mu_shift=0.028 sigma=43.411
Dihedral - 177.8 deg
Dihedral: 3-13-4-17
AA mu_shift=0.000 sigma=37.659
CG mu_shift=0.680 sigma=40.410
Dihedral - 177.9 deg
Dihedral: 3-13-12-14
AA mu_shift=0.000 sigma=12.509
CG mu_shift=-0.319 sigma=14.131
Dihedral - -179.6 deg
Dihedral: 4-13-3-10
AA mu_shift=0.000 sigma=36.995
CG mu_shift=-0.397 sigma=40.427
Dihedral - -178.1 deg
Dihedral: 4-13-3-12
AA mu_shift=0.000 sigma=37.465
CG mu_shift=0.028 sigma=40.724
Dihedral - 1.8 deg
Dihedral: 4-13-12-14
AA mu_shift=0.000 sigma=14.625
CG mu_shift=-0.421 sigma=15.253
Dihedral - 0.8 deg
Dihedral: 4-15-16-18
AA mu_shift=0.000 sigma=39.996
CG mu_shift=-11.482 sigma=55.070
Dihedral - -133.8 deg
Dihedral: 4-17-1-5
AA mu_shift=0.000 sigma=75.433
CG mu_shift=-9.358 sigma=94.573
Dihedral - -19.3 deg
Dihedral: 4-17-1-7
AA mu_shift=0.000 sigma=75.328
CG mu_shift=-4.333 sigma=86.728
Dihedral - -10.0 deg
Dihedral: 4-17-16-18
AA mu_shift=0.000 sigma=39.315
CG mu_shift=-8.445 sigma=83.349
Dihedral - 104.2 deg
Dihedral: 5-1-17-16
AA mu_shift=0.000 sigma=32.986
CG mu_shift=5.688 sigma=37.161
Dihedral - 157.8 deg
Dihedral: 6-5-1-17
AA mu_shift=0.000 sigma=43.511
CG mu_shift=-8.530 sigma=45.072
Dihedral - 16.9 deg
Dihedral: 6-7-1-17
AA mu_shift=0.000 sigma=12.451
CG mu_shift=-0.487 sigma=14.161
Dihedral - 179.8 deg
Dihedral: 6-7-2-9
AA mu_shift=0.000 sigma=12.933
CG mu_shift=-1.071 sigma=18.564
Dihedral - -179.8 deg
Dihedral: 6-7-2-10
AA mu_shift=0.000 sigma=12.678
CG mu_shift=0.363 sigma=14.514
Dihedral - -179.5 deg
Dihedral: 7-1-5-8
AA mu_shift=0.000 sigma=50.281
CG mu_shift=-4.819 sigma=42.388
Dihedral - -153.1 deg
Dihedral: 7-1-17-16
AA mu_shift=0.000 sigma=8.805
CG mu_shift=-0.315 sigma=11.535
Dihedral - 168.7 deg
Dihedral: 7-2-9-10
AA mu_shift=0.000 sigma=18.054
CG mu_shift=1.545 sigma=18.244
Dihedral - 0.3 deg
Dihedral: 7-2-9-11
AA mu_shift=0.000 sigma=24.697
CG mu_shift=-0.646 sigma=23.915
Dihedral - 2.0 deg
Dihedral: 7-2-10-9
AA mu_shift=0.000 sigma=8.701
CG mu_shift=0.705 sigma=8.714
Dihedral - -179.8 deg
Dihedral: 7-6-5-8
AA mu_shift=0.000 sigma=76.166
CG mu_shift=21.404 sigma=60.760
Dihedral - 131.8 deg
Dihedral: 8-5-1-17
AA mu_shift=0.000 sigma=73.071
CG mu_shift=-24.964 sigma=60.422
Dihedral - -127.9 deg
Dihedral: 9-10-3-12
AA mu_shift=0.000 sigma=19.135
CG mu_shift=0.221 sigma=22.855
Dihedral - 179.9 deg
Dihedral: 9-10-3-13
AA mu_shift=0.000 sigma=10.941
CG mu_shift=-0.661 sigma=12.203
Dihedral - -180.0 deg
Dihedral: 10-2-9-11
AA mu_shift=0.000 sigma=15.210
CG mu_shift=-2.138 sigma=17.105
Dihedral - 1.7 deg
Dihedral: 10-3-12-13
AA mu_shift=0.000 sigma=18.490
CG mu_shift=-0.917 sigma=19.867
Dihedral - 0.1 deg
Dihedral: 10-3-12-14
AA mu_shift=0.000 sigma=25.943
CG mu_shift=-1.397 sigma=25.561
Dihedral - 0.6 deg
Dihedral: 10-3-13-12
AA mu_shift=0.000 sigma=7.155
CG mu_shift=0.279 sigma=7.477
Dihedral - 180.0 deg
Dihedral: 12-13-4-15
AA mu_shift=0.000 sigma=24.588
CG mu_shift=0.145 sigma=26.172
Dihedral - 179.3 deg
Dihedral: 12-13-4-17
AA mu_shift=0.000 sigma=8.529
CG mu_shift=0.592 sigma=10.839
Dihedral - 179.6 deg
Dihedral: 13-3-12-14
AA mu_shift=0.000 sigma=15.538
CG mu_shift=-0.451 sigma=17.253
Dihedral - 0.5 deg
Dihedral: 13-4-15-16
AA mu_shift=0.000 sigma=28.587
CG mu_shift=0.315 sigma=28.946
Dihedral - -10.3 deg
Dihedral: 13-4-17-16
AA mu_shift=0.000 sigma=10.305
CG mu_shift=0.102 sigma=11.567
Dihedral - -168.8 deg

### LIB

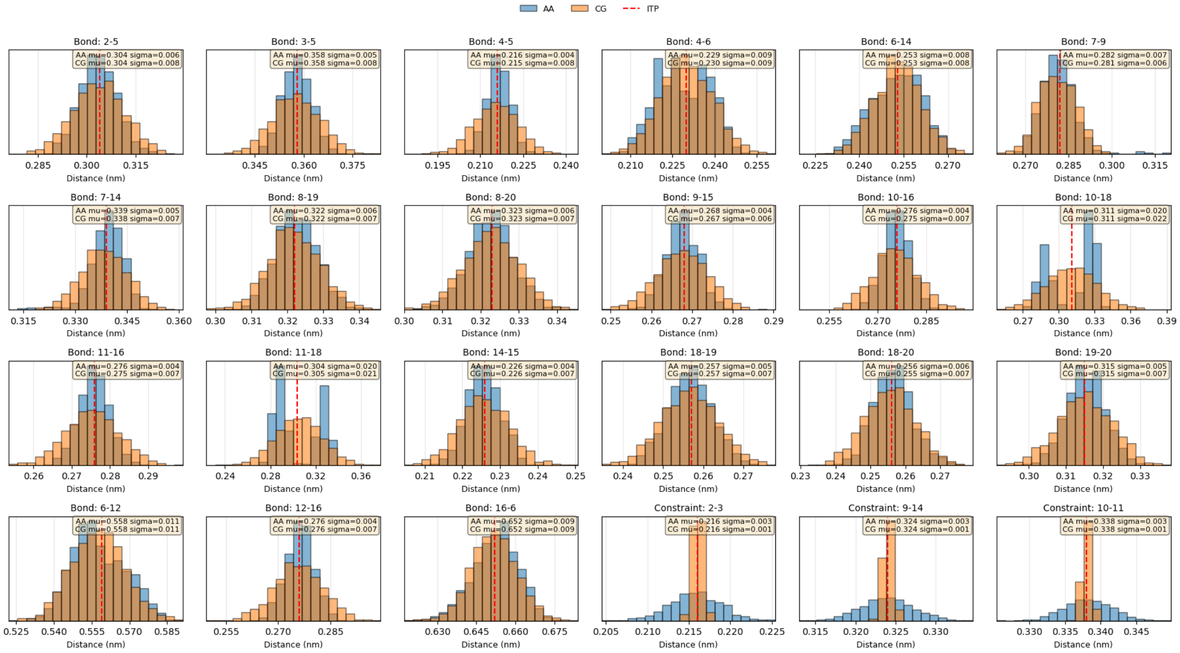
AA
CG
ITP
Bond: 2-5
Bond: 3-5
Bond: 4-5
Bond: 4-6
Bond: 6-14
Bond: 7-9
Bond: 7-14
Bond: 8-19
Bond: 8-20
Bond: 9-15
Bond: 10-16
Bond: 10-18
Bond: 11-16
Bond: 11-18
Bond: 14-15
Bond: 18-19
Bond: 18-20
Bond: 19-20
Bond: 6-12
Bond: 12-16
Bond: 16-6
Constraint: 2-3
Constraint: 9-14
Constraint: 10-11
Distance (nm)

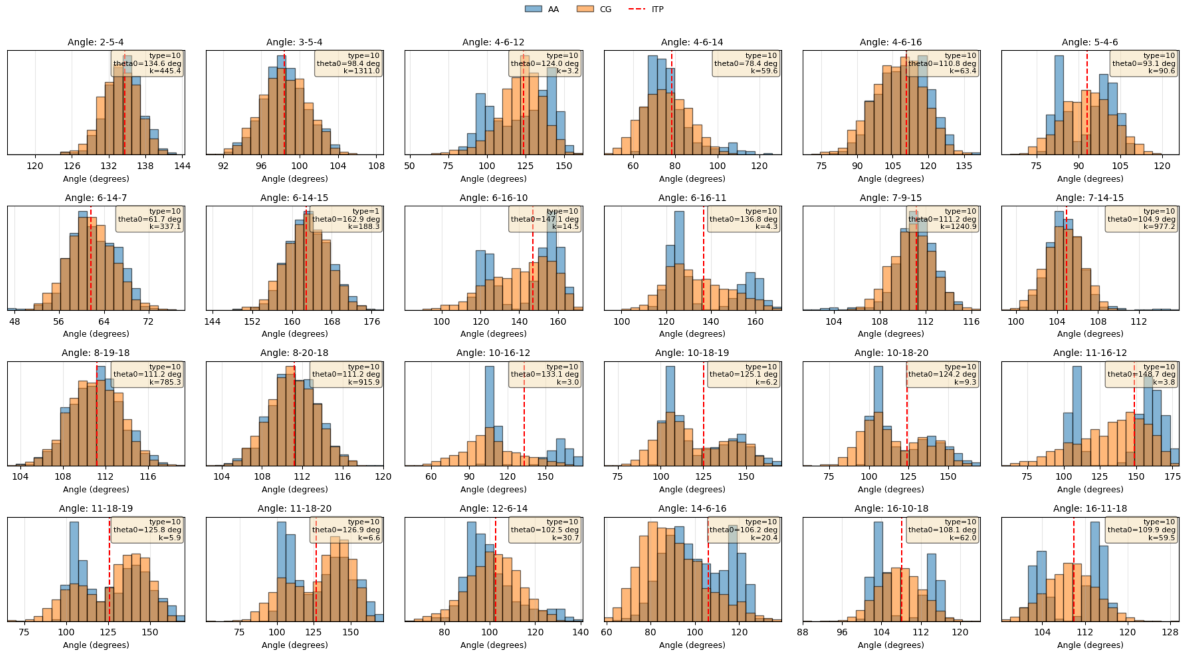
AA
CG
ITP
Angle: 2-5-4
Angle: 3-5-4
Angle: 4-6-12
Angle: 4-6-14
Angle: 4-6-16
Angle: 5-4-6
Angle: 6-14-7
Angle: 6-14-15
Angle: 6-16-10
Angle: 6-16-11
Angle: 7-9-15
Angle: 7-14-15
Angle: 8-19-18
Angle: 8-20-18
Angle: 10-16-12
Angle: 10-18-19
Angle: 10-18-20
Angle: 11-16-12
Angle: 11-18-19
Angle: 11-18-20
Angle: 12-6-14
Angle: 14-6-16
Angle: 16-10-18
Angle: 16-11-18
Angle (degrees)

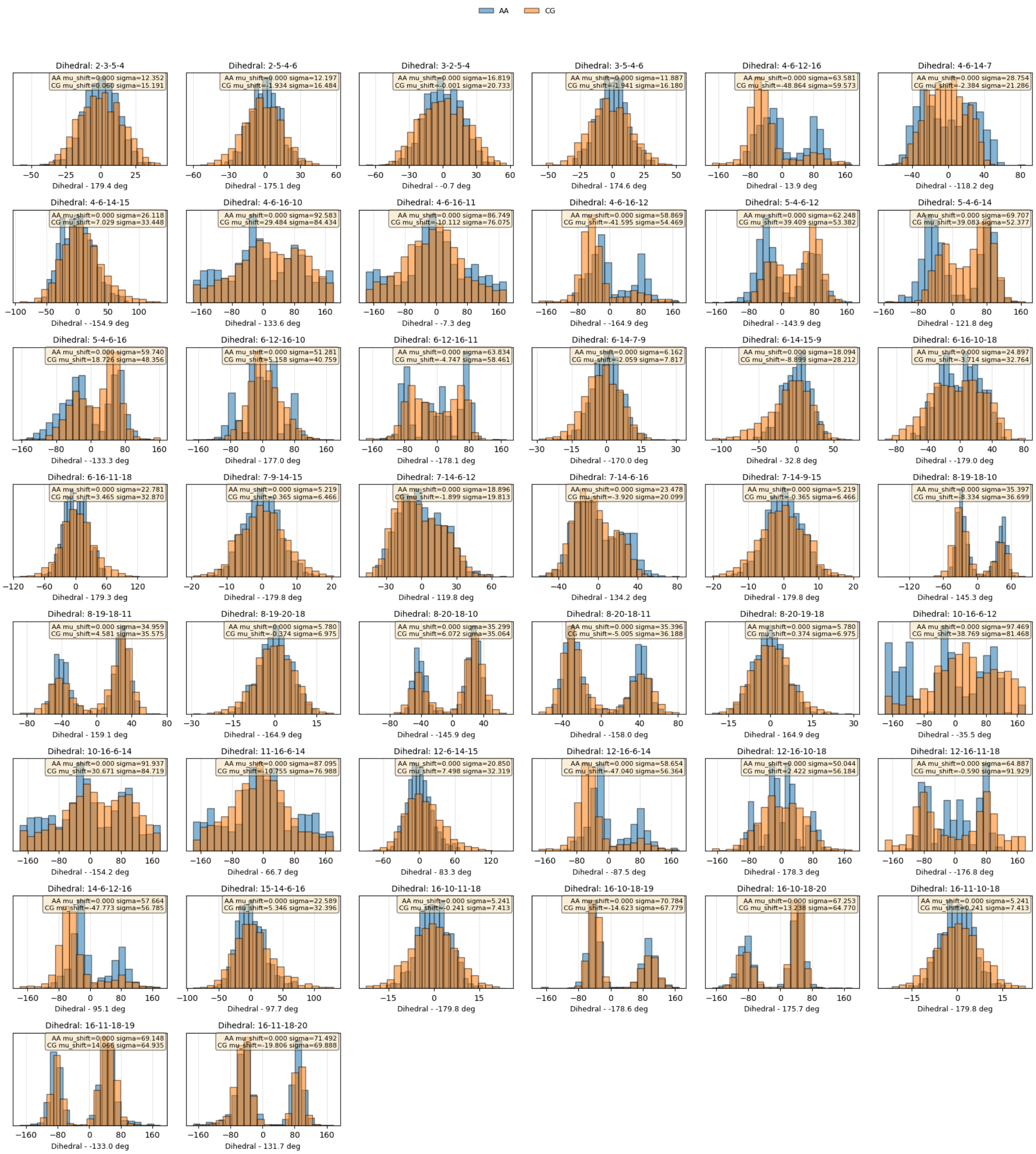
AA
CG
Dihedral: 2-3-5-4
AA mu_shift=0.000 sigma=12.352
CG mu_shift=0.060 sigma=15.191
Dihedral - 179.4 deg
Dihedral: 2-5-4-6
AA mu_shift=0.000 sigma=12.197
CG mu_shift=-1.934 sigma=16.484
Dihedral - 175.1 deg
Dihedral: 3-2-5-4
AA mu_shift=0.000 sigma=16.819
CG mu_shift=-0.001 sigma=20.733
Dihedral - -0.7 deg
Dihedral: 3-5-4-6
AA mu_shift=0.000 sigma=11.887
CG mu_shift=-1.941 sigma=16.180
Dihedral - 174.6 deg
Dihedral: 4-6-12-16
AA mu_shift=0.000 sigma=63.581
CG mu_shift=-48.864 sigma=59.573
Dihedral - 13.9 deg
Dihedral: 4-6-14-7
AA mu_shift=0.000 sigma=28.754
CG mu_shift=-2.384 sigma=21.286
Dihedral - -118.2 deg
Dihedral: 4-6-14-15
AA mu_shift=0.000 sigma=26.118
CG mu_shift=7.029 sigma=33.448
Dihedral - -154.9 deg
Dihedral: 4-6-16-10
AA mu_shift=0.000 sigma=92.583
CG mu_shift=29.484 sigma=84.434
Dihedral - 133.6 deg
Dihedral: 4-6-16-11
AA mu_shift=0.000 sigma=86.749
CG mu_shift=-10.112 sigma=76.075
Dihedral - -7.3 deg
Dihedral: 4-6-16-12
AA mu_shift=0.000 sigma=58.869
CG mu_shift=-41.595 sigma=54.469
Dihedral - -164.9 deg
Dihedral: 5-4-6-12
AA mu_shift=0.000 sigma=62.248
CG mu_shift=39.409 sigma=53.382
Dihedral - -143.9 deg
Dihedral: 5-4-6-14
AA mu_shift=0.000 sigma=69.707
CG mu_shift=39.083 sigma=52.377
Dihedral - 121.8 deg
Dihedral: 5-4-6-16
AA mu_shift=0.000 sigma=59.740
CG mu_shift=18.726 sigma=48.356
Dihedral - -133.3 deg
Dihedral: 6-12-16-10
AA mu_shift=0.000 sigma=51.281
CG mu_shift=5.158 sigma=40.759
Dihedral - 177.0 deg
Dihedral: 6-12-16-11
AA mu_shift=0.000 sigma=63.834
CG mu_shift=-4.747 sigma=58.461
Dihedral - -178.1 deg
Dihedral: 6-14-7-9
AA mu_shift=0.000 sigma=6.162
CG mu_shift=-2.059 sigma=7.817
Dihedral - -170.0 deg
Dihedral: 6-14-15-9
AA mu_shift=0.000 sigma=18.094
CG mu_shift=-8.899 sigma=28.212
Dihedral - 32.8 deg
Dihedral: 6-16-10-18
AA mu_shift=0.000 sigma=24.897
CG mu_shift=-3.714 sigma=32.764
Dihedral - -179.0 deg
Dihedral: 6-16-11-18
AA mu_shift=0.000 sigma=22.781
CG mu_shift=3.465 sigma=32.870
Dihedral - 179.3 deg
Dihedral: 7-9-14-15
AA mu_shift=0.000 sigma=5.219
CG mu_shift=0.365 sigma=6.466
Dihedral - -179.8 deg
Dihedral: 7-14-6-12
AA mu_shift=0.000 sigma=18.896
CG mu_shift=-1.899 sigma=19.813
Dihedral - 119.8 deg
Dihedral: 7-14-6-16
AA mu_shift=0.000 sigma=23.478
CG mu_shift=-3.920 sigma=20.099
Dihedral - 134.2 deg
Dihedral: 7-14-9-15
AA mu_shift=0.000 sigma=5.219
CG mu_shift=-0.365 sigma=6.466
Dihedral - 179.8 deg
Dihedral: 8-19-18-10
AA mu_shift=0.000 sigma=35.397
CG mu_shift=-8.334 sigma=36.699
Dihedral - 145.3 deg
Dihedral: 8-19-18-11
AA mu_shift=0.000 sigma=34.959
CG mu_shift=4.581 sigma=35.575
Dihedral - 159.1 deg
Dihedral: 8-19-20-18
AA mu_shift=0.000 sigma=5.780
CG mu_shift=-0.374 sigma=6.975
Dihedral - -164.9 deg
Dihedral: 8-20-18-10
AA mu_shift=0.000 sigma=35.299
CG mu_shift=6.072 sigma=35.064
Dihedral - -145.9 deg
Dihedral: 8-20-18-11
AA mu_shift=0.000 sigma=35.396
CG mu_shift=-5.005 sigma=36.188
Dihedral - -158.0 deg
Dihedral: 8-20-19-18
AA mu_shift=0.000 sigma=5.780
CG mu_shift=0.374 sigma=6.975
Dihedral - 164.9 deg
Dihedral: 10-16-6-12
AA mu_shift=0.000 sigma=97.469
CG mu_shift=38.769 sigma=81.468
Dihedral - -35.5 deg
Dihedral: 10-16-6-14
AA mu_shift=0.000 sigma=91.937
CG mu_shift=30.671 sigma=84.719
Dihedral - -154.2 deg
Dihedral: 11-16-6-14
AA mu_shift=0.000 sigma=87.095
CG mu_shift=-10.755 sigma=76.988
Dihedral - 66.7 deg
Dihedral: 12-6-14-15
AA mu_shift=0.000 sigma=20.850
CG mu_shift=7.498 sigma=32.319
Dihedral - 83.3 deg
Dihedral: 12-16-6-14
AA mu_shift=0.000 sigma=58.654
CG mu_shift=-47.040 sigma=56.364
Dihedral - -87.5 deg
Dihedral: 12-16-10-18
AA mu_shift=0.000 sigma=50.044
CG mu_shift=2.422 sigma=56.184
Dihedral - 178.3 deg
Dihedral: 12-16-11-18
AA mu_shift=0.000 sigma=64.887
CG mu_shift=-0.590 sigma=91.929
Dihedral - -176.8 deg
Dihedral: 14-6-12-16
AA mu_shift=0.000 sigma=57.664
CG mu_shift=-47.773 sigma=56.785
Dihedral - 95.1 deg
Dihedral: 15-14-6-16
AA mu_shift=0.000 sigma=22.589
CG mu_shift=5.346 sigma=32.396
Dihedral - 97.7 deg
Dihedral: 16-10-11-18
AA mu_shift=0.000 sigma=5.241
CG mu_shift=-0.241 sigma=7.413
Dihedral - -179.8 deg
Dihedral: 16-10-18-19
AA mu_shift=0.000 sigma=70.784
CG mu_shift=-14.623 sigma=67.779
Dihedral - -178.6 deg
Dihedral: 16-10-18-20
AA mu_shift=0.000 sigma=67.253
CG mu_shift=13.238 sigma=64.770
Dihedral - 175.7 deg
Dihedral: 16-11-10-18
AA mu_shift=0.000 sigma=5.241
CG mu_shift=0.241 sigma=7.413
Dihedral - 179.8 deg
Dihedral: 16-11-18-19
AA mu_shift=0.000 sigma=69.148
CG mu_shift=14.068 sigma=64.935
Dihedral - -133.0 deg
Dihedral: 16-11-18-20
AA mu_shift=0.000 sigma=71.492
CG mu_shift=-19.806 sigma=69.888
Dihedral - 131.7 deg

### THC

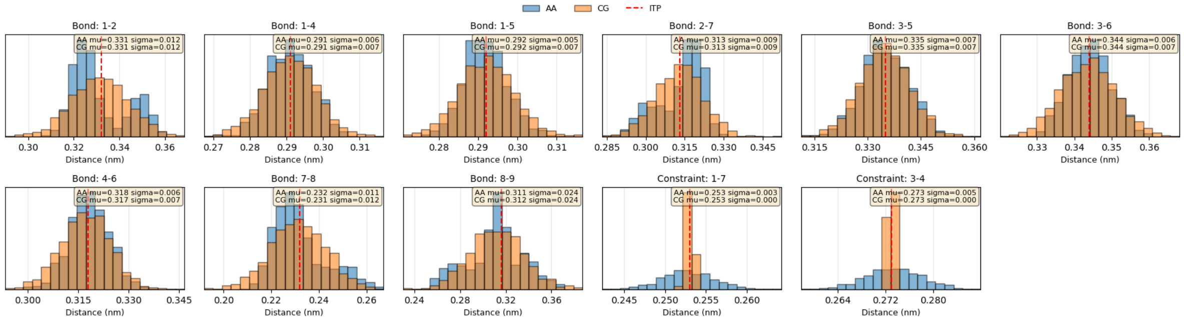
AA
CG
ITP
Bond: 1-2
AA mu=0.331 sigma=0.012
CG mu=0.331 sigma=0.012
Bond: 1-4
AA mu=0.291 sigma=0.006
CG mu=0.291 sigma=0.007
Bond: 1-5
AA mu=0.292 sigma=0.005
CG mu=0.292 sigma=0.007
Bond: 2-7
AA mu=0.313 sigma=0.009
CG mu=0.313 sigma=0.009
Bond: 3-5
AA mu=0.335 sigma=0.007
CG mu=0.335 sigma=0.007
Bond: 3-6
AA mu=0.344 sigma=0.006
CG mu=0.344 sigma=0.007
Bond: 4-6
AA mu=0.318 sigma=0.006
CG mu=0.317 sigma=0.007
Bond: 7-8
AA mu=0.232 sigma=0.011
CG mu=0.231 sigma=0.012
Bond: 8-9
AA mu=0.311 sigma=0.024
CG mu=0.312 sigma=0.024
Constraint: 1-7
AA mu=0.253 sigma=0.003
CG mu=0.253 sigma=0.000
Constraint: 3-4
AA mu=0.273 sigma=0.005
CG mu=0.273 sigma=0.000
Distance (nm)

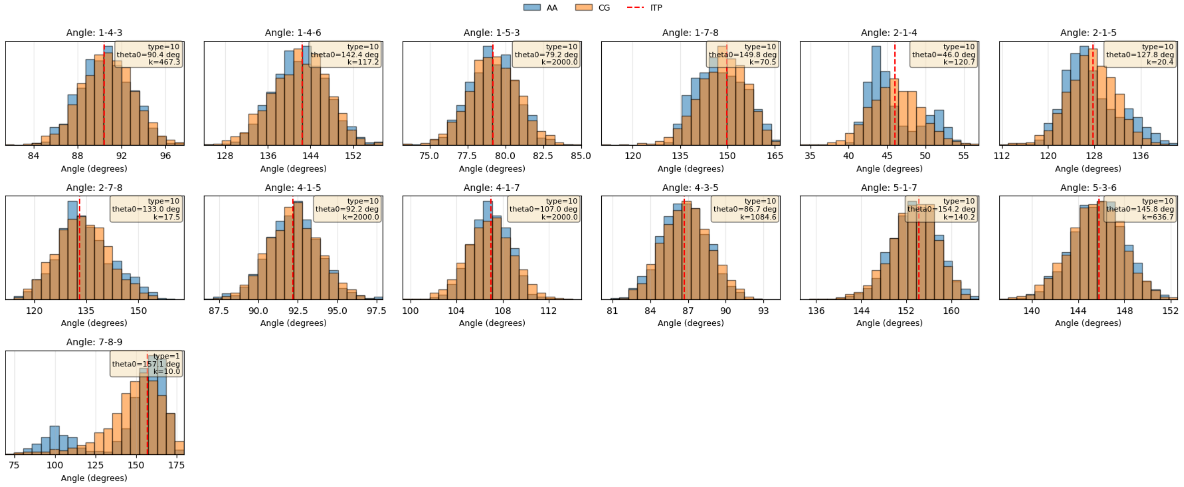
AA
CG
ITP
Angle: 1-4-3
type=10
theta0=90.4 deg
k=467.3
Angle: 1-4-6
type=10
theta0=142.4 deg
k=117.2
Angle: 1-5-3
type=10
theta0=79.2 deg
k=2000.0
Angle: 1-7-8
type=10
theta0=149.8 deg
k=70.5
Angle: 2-1-4
type=10
theta0=46.0 deg
k=120.7
Angle: 2-1-5
type=10
theta0=127.8 deg
k=20.4
Angle: 2-7-8
type=10
theta0=133.0 deg
k=17.5
Angle: 4-1-5
type=10
theta0=92.2 deg
k=2000.0
Angle: 4-1-7
type=10
theta0=107.0 deg
k=2000.0
Angle: 4-3-5
type=10
theta0=86.7 deg
k=1084.6
Angle: 5-1-7
type=10
theta0=154.2 deg
k=140.2
Angle: 5-3-6
type=10
theta0=145.8 deg
k=636.7
Angle: 7-8-9
type=1
theta0=157.1 deg
k=10.0
Angle (degrees)

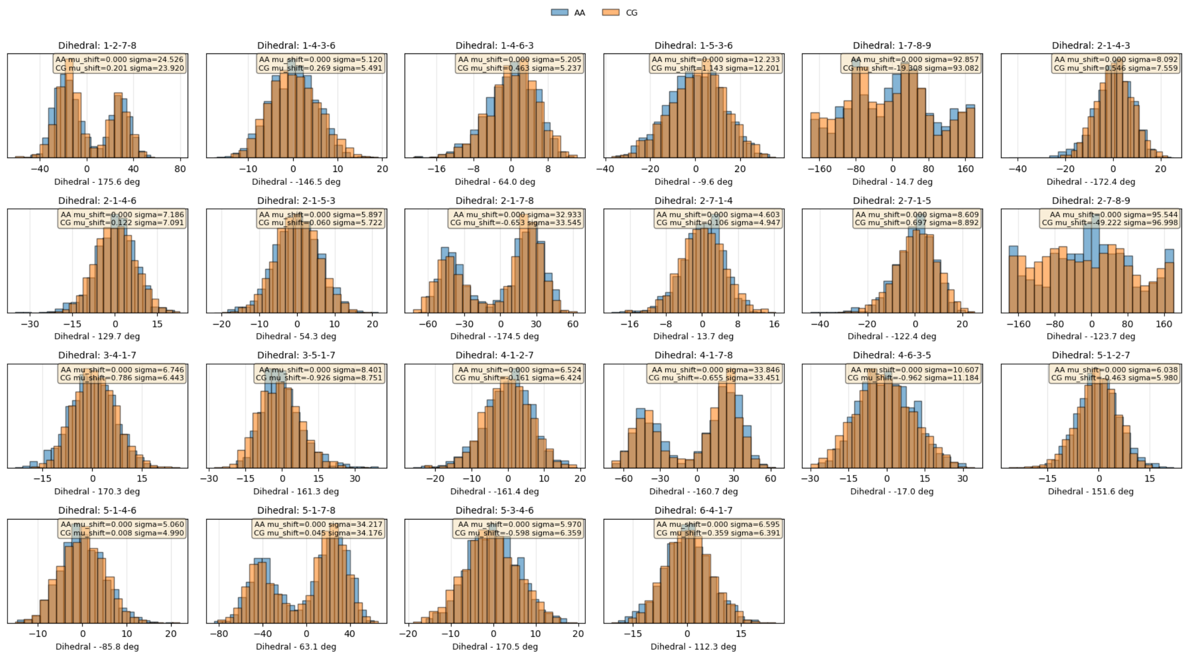
